# Deep profiling of lupus nephritis kidneys reveals dynamic changes in myeloid cells associated with disease progression

**DOI:** 10.64898/2025.12.17.694954

**Authors:** Paul J. Hoover, Thomas M. Eisenhaure, Jeffrey Hodgin, William Apruzzese, Joseph Mears, Michael Peters, Tony Jones, Sujal I. Shah, Haniya Kamal, Rollin Leavitt, Shaun W. Jackson, Patrick Danaher, Deepak A. Rao, Qian Xiao, Siddarth Gurajala, Andrea Fava, Celine C. Berthier, Alice Horisberger, Peter M. Izmirly, H. Michael Belmont, Robert Clancy, Richard Furie, Joel M. Guthridge, Maria Dall’Era, David Wofsy, Diane L. Kamen, Kenneth C. Kalunian, Maureen A. McMahon, Jennifer Grossman, Fernanda Payan-Schober, David A. Hildeman, E. Steve Woodle, Chaim Putterman, Matthias Kretzler, Soumya Raychaudhuri, Judith A. James, Jennifer H. Anolik, Michelle A. Petri, Jill P. Buyon, the Accelerating Medicines Partnership (AMP) RA/SLE Network, Betty Diamond, Anne Davidson, Nir Hacohen, Arnon Arazi

## Abstract

**Objectives:** Lupus nephritis (LN) is a common, potentially fatal manifestation of systemic lupus erythematosus. We aim to gain new insights into the immune responses underlying LN and their relation to the histologic heterogeneity observed in this disease, focusing on myeloid cells.

**Methods:** Single-cell RNA-sequencing (scRNA-seq) was used to profile dissociated kidney samples from 156 LN patients and 30 healthy individuals. Spatial transcriptomics (ST), utilizing a gene panel designed to capture all myeloid subsets identified in the scRNA-seq data, was applied to kidney samples acquired from 6 LN patients and 2 healthy controls.

**Results:** We generated a full catalog of the myeloid subsets found in LN kidneys. Our analyses indicated that an increase in irreversible tissue damage, as measured by the NIH chronicity index (CI), is associated with a gradual switch of the local immune response from one dominated by monocytes and macrophages to one featuring expanded CD4^+^ T, GZMK^+^CD8^+^ T, B and dendritic cells, with a parallel decrease in the interferon response. In proliferative/mixed LN only, the degree of active inflammation correlates with expansion of disease-specific macrophage (DMac) subsets, which later contract as the CI increases. Trajectory analysis of the scRNA-seq data suggested that DMacs arise from both infiltrating monocytes and tissue-resident macrophages; this was supported by the ST data, as well as cell cultures. DMacs are indicated to interact with parietal epithelial cells, promoting the development of glomerulosclerosis.

**Conclusions:** We suggest a detailed picture of the changes in the kidney immune mechanisms in LN as this disease progresses.

## INTRODUCTION

Lupus nephritis (LN) is a common and severe manifestation of systemic lupus erythematosus (SLE), occurring in ∼50% of patients and leading to end stage renal disease in ∼10-45% of cases within 15 years of diagnosis [1]. Unfortunately, the response rate of LN patients to therapy remains below 50% even after new drugs have been approved for its treatment in recent years [2, 3]. Thus, there is an urgent need to gain better understanding of the immune mechanisms driving LN, in particular those operating in the kidney, to enable the development of more effective and specific drugs.

LN kidney histopathology is traditionally described by the International Society of Nephrology (ISN) classification and two composite indices - the NIH activity and chronicity indices (AI and CI, respectively) [4]. The AI reflects the degree of active kidney inflammation, while the CI roughly represents the extent of irreversible kidney damage; patients with high CI are generally less responsive to immunotherapy [5, 6].

Immune cells of myeloid origin, in particular macrophages and dendritic cells (DCs), demonstrate high phenotypic and functional complexity in LN kidneys [7]. Compared to mouse models of LN, the current knowledge regarding macrophages and DCs in human patients is limited. The importance of myeloid cells has been demonstrated in studies showing a correlation between their presence in LN kidneys and disease severity, as well as the likelihood of disease progression or treatment outcome [8–10]. We have previously used single-cell RNA-sequencing (scRNA-seq) to profile kidneys from a small cohort of LN patients, and showed the presence of both proinflammatory and alternatively-activated macrophages [11]. However, a full catalog of the different subsets of myeloid cells in LN kidneys, their relation to disease heterogeneity as represented by the ISN class, AI and CI, their exact position in the kidney, their likely origin and their role in disease progression are largely unknown.

To address these questions, we generated and analyzed a large-scale scRNA-seq dataset of dissociated kidney biopsies collected from a diverse cohort of LN patients and healthy living donors (LD). We complemented these data with spatial transcriptomics (ST) of LN kidney samples, utilizing a gene panel customized to capture all myeloid cell subsets found in the scRNA-seq data. While our focus is myeloid cells, we also explored their balance with other immune cells and its relation to disease progression. In addition, we investigated the interactions of myeloid cells with the tissue itself, focusing on parietal epithelial cells (PECs), and the way such interactions may promote disease processes.

## METHODS

See additional details in the Supplementary Information file.

### Patients

Kidney biopsies were obtained from 156 LN patients to confirm suspected LN de novo, relapse of disease or activity not responding to treatment, as well as 30 LD controls (Supplemental Table 1). The study protocol was approved by the institutional review boards and ethics committees of participating sites in adherence with the Declaration of Helsinki and all informed consent was obtained from all participants.

### Patient and public involvement

Patients and the public were not involved in the design, conduct, reporting, and dissemination plans of this manuscript.

### Single-cell data

scRNA-seq was performed on biopsies collected from our entire cohort, using the 10x Genomics Chromium platform (3’ V3 kit). Sequencing was done using Illumina NovaSeq S4.

### Spatial transcriptomics

ST of kidney biopsies from 6 LN patients and 2 healthy controls was performed using 10x Xenium.

### Data analysis

Clustering of scRNA-seq data was performed using Seurat v4. Trajectory analysis was done using PAGA [12]. Additional analyses were performed using designated code.

## RESULTS

### Clustering identifies disease-specific subsets of monocytes and macrophages

The scRNA-seq data generated from our cohort contained 23,819 myeloid cells, which constituted 32% of kidney immune cells (Supplementary Fig. 1a,b. A detailed analysis of T and NK cells is included elsewhere (Al Souz et al, manuscript under revision)). High-resolution clustering of kidney myeloid cells identified 24 cell subsets (Fig. 1a). The labeling of these clusters was based on their relative frequency in LN patients and LD controls (Fig. 1b; Supplementary Fig. 1c), the expression of canonical markers (Fig. 1c; Supplementary Table 2), and cell mapping to published reference scRNA-seq datasets characterizing the contents of kidneys and blood obtained from healthy individuals [13, 14] (Fig. 1d; Supplementary Fig. 1d). We identified clusters of classical, nonclassical and intermediate monocytes, DCs (cDC1, cDC2, CCR7^+^ DCs and pDCs), Lyve1^+^ and Lyve1^−^ tissue-resident macrophages (RMs) and mast cells. The clusters commonly found in LD controls – monocytes, RMs and cDC2 – all demonstrated elevated expression of interferon stimulated genes (ISGs) in LN patients, as expected (Supplementary Fig. 1e).

**Figure 1.**
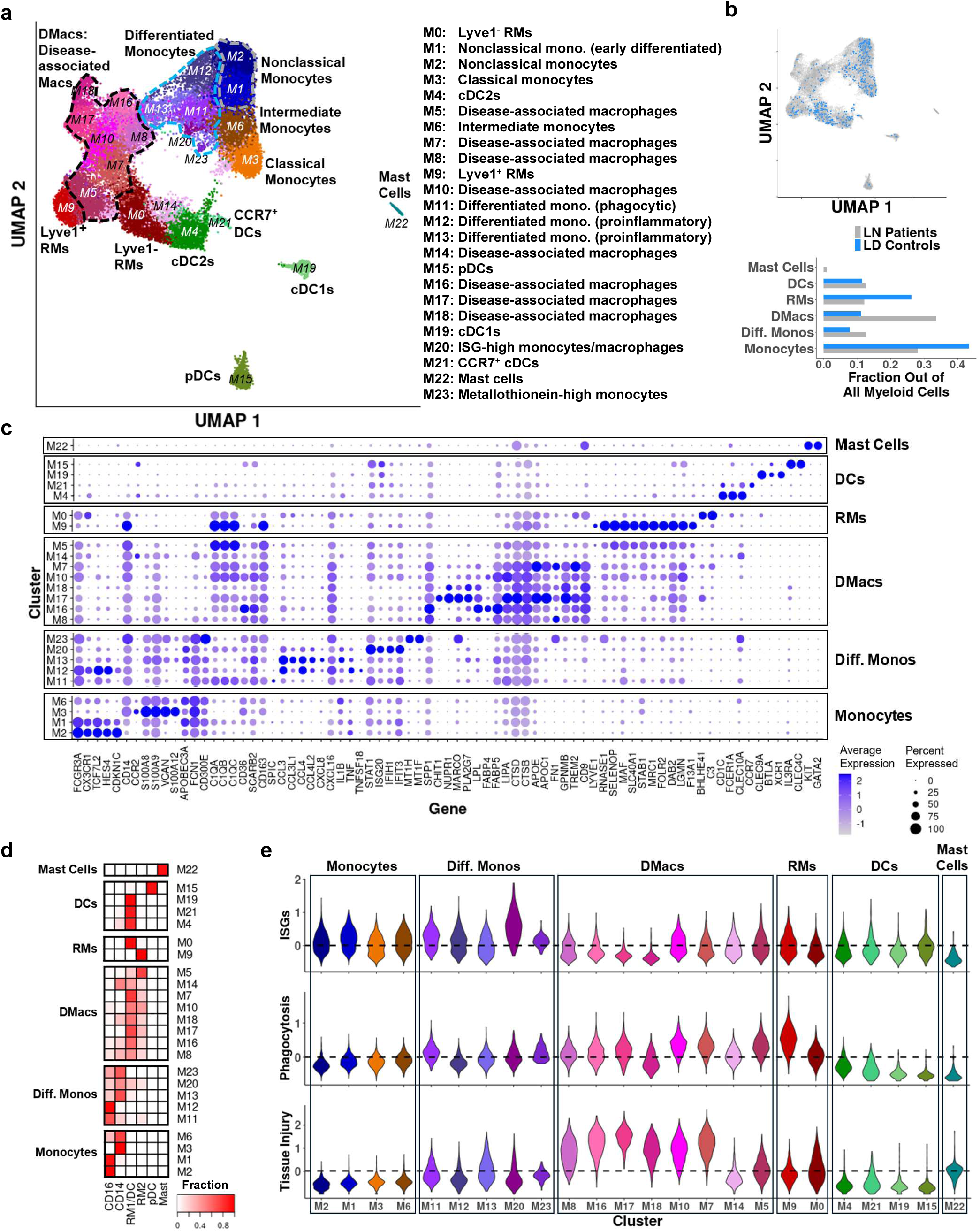
Myeloid cell clusters in LN kidneys. (a) Myeloid cell clusters and their putative labels. (b) The distribution of cells extracted from LN patients and LD controls in the UMAP plane (top), and the fraction of each class of clusters in LN patients and LD controls (bottom). (c) The expression of selected genes across the myeloid cell clusters. (d) The fraction of cells mapped to each reference class in Stewart et. al, 2019, specified for each myeloid cell cluster. (e) The distribution of three composite scores, computed per cell as the average of scaled expression of a gene set of interest, across myeloid cell clusters. Only cells from LN patients were taken into account. The dashed line represents the average across all cells from LN patients, regardless of cluster. Top: the “interferon score”, computed over a set of ISGs. Middle: a composite score computed based on genes associated with phagocytosis. Bottom, a composite score computed using genes associated with tissue injury.

Clusters M11, M12, M13, M20 and M23, which were largely missing in LD controls, were exclusively mapped to monocytes, indicating that they were unlikely to originate from RMs. Cluster M11, suggested to be a mixture of classical and nonclassical monocytes, demonstrated relatively high levels of both ISGs and genes associated with phagocytosis (Fig. 1e; Supplementary Fig. 1f; Supplementary Table 3). Cluster M12, composed mostly of nonclassical monocytes, demonstrated lower levels of phagocytosis-related genes compared to M11, and upregulation of several chemokines and proinflammatory cytokines. Cluster M13, consisting of both classical and nonclassical monocytes, showed a similar transcriptional pattern. Cluster M20 demonstrated particularly high expression of ISGs; while this cluster was predominantly composed of monocytes, it did include some cells likely originating from RMs and DCs. Cluster M23 was characterized by the upregulation of several metallothionein genes.

Cell mapping indicated that clusters M5, M7, M8, M10, M14, M16, M17 and M18 were a mixture of cells derived from RMs and monocytes. These clusters were highly-expanded in LN patients compared to LD controls, and were thus labeled “disease-associated macrophages” (DMacs). Cluster M5 presented an expression signature most similar to the Lyve1^+^ RMs (cluster M9), and likely represents early differentiation from these cells. This signature was also observed to a lesser extent in clusters M7 and M10. These two clusters, as well as clusters M8, M16, M17 and M18, featured high expression levels of injury-associated genes. Compared to the other DMacs, cluster M18 demonstrated a relatively low expression of phagocytosis-associated genes. Clusters M18 and M17 showed a relatively low interferon response; the latter cluster, together with clusters M7, and M16, was characterized by the upregulation of extracellular matrix (ECM) regulators. Cluster M17, and to a lesser degree cluster M18, featured the upregulation of several genes associated with lipid metabolism.

### The CI is correlated with relative contraction of monocytes and macrophages, and relative expansion of B, memory CD4^+^ and CD8^+^GZMK^+^ T cells

Clustering of LN patients based on immune cell composition, considering both myeloid and lymphoid cell subsets, did not identify distinct groups of LN patients, but rather pointed to a continuum of cell mixtures across the cohort. We therefore used a continuous approach - principal component analysis (PCA) - to characterize this heterogeneity. PC1, capturing the main source (33%) of the total variance, represented the balance between lymphocytes - in particular B cells, GZMK^+^CD8^+^ T cells and memory CD4^+^ T cells - and myeloid cells, specifically monocytes and DMacs (Fig. 2a). For both proliferative/mixed and pure membranous LN, PC1 was significantly correlated with the CI, such that an increase in the CI was associated with a higher ratio between the above lymphoid and myeloid subsets (Fig. 2b; Supplementary Fig. 2a-e). These observations were not driven by age, race, ethnicity, sex, treatment, or by whether the biopsy was acquired at first onset or at relapse of disease (Supplementary Table 4).

**Figure 2.**
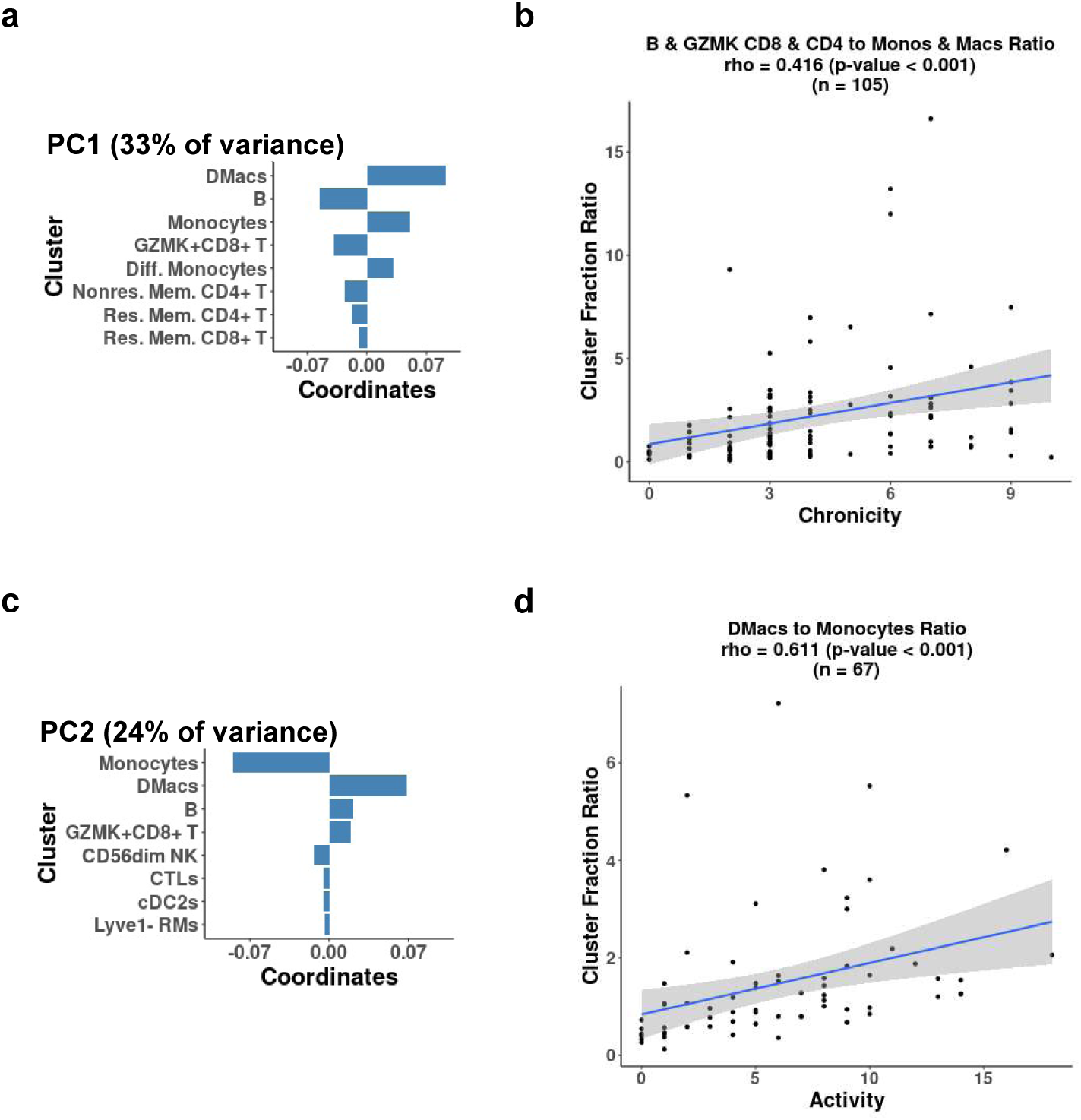
The relation between the global frequencies of immune cell subsets and histopathological features. (a) The top contributing clusters to PC1, in PCA analysis of immune cluster fractions (out of all immune cells), considering all LN patients with at least 20 immune cells; the reported results were found to be robust with regard to the choice of this threshold, within the range 10-100 cells. The percentage of total variance explained by PC1 is specified in parentheses. (b) The relation between the NIH chronicity index (CI) and the ratio between the relative frequencies of B cells, GZMK^+^CD8^+^ T cells and memory CD4^+^ T cells, and that of monocytes and macrophages. Reported are the Spearman correlation and its associated p-value, as well as the number of samples used in their calculation. (c) Same as panel (a), for PC2. (d) The relation between the NIH activity index (AI) and the ratio between selected clusters corresponding to differentiated macrophages and undifferentiated monocytes. In panels (b) and (d), only LN patients with at least 20 cells in both numerator and denominator of the calculated ratio were considered; we found the results to be robust with regard to the exact choice of this threshold, within the range 10-100 cells.

Since the CI represents irreversible kidney damage and is expected to generally increase with the duration of active disease [15–17], the observed correlations suggest disease progression is accompanied by a gradual transition in the kidney immune response, initially dominated by monocytes and macrophages, to one increasingly driven by lymphocytes. Accordingly, both PC1 and the ratio between these immune cell subsets showed significant correlations with the patients’ age at the time of biopsy (Supplementary Fig. 2f-g).

### In proliferative/mixed LN, the AI is correlated with relative expansion of DMacs, while increased CI is correlated with a decrease in DMacs and concurrent expansion of DCs

PC2 reflected the balance between monocytes and DMacs (Fig. 2c). We found that in proliferative/mixed LN, but not in pure membranous disease, PC2 showed a significant correlation with the AI. Specifically, higher values of the AI corresponded to a relative expansion of DMacs and a relative contraction of monocytes (Fig. 2d; Supplementary Fig. 3a-c), suggesting differentiation of monocytes into DMacs.

Next, we focused our analysis on myeloid cells. We examined the relation between cluster frequencies and the CI and AI, analyzing separately patients with proliferative or pure membranous disease. Patients with mixed proliferative/membranous disease (classes III/V and IV/V) were included in the proliferative group, as they demonstrated similar patterns of immune cell composition and gene expression. Within each ISN class, patients were further stratified by their CI into high (CI≥5) and low (CI≤2) groups (Figs. 3a, 4a). Since lower CI generally reflects earlier disease stages, this enabled comparing early and more advanced disease. For patients with proliferative/mixed LN, we further required the AI to be ≥3, to avoid including patients with “burnt-out” LN [18, 19], where active inflammation subsides following substantial kidney damage, since this late disease stage likely involves different immune mechanisms. For patients with pure membranous disease we did not apply an AI threshold, since their AI was uniformly low, as expected. The results reported below were consistent regardless of the specific thresholds used for the AI and CI, demonstrating their robustness (Supplementary Figs. 4-8).

**Figure 3.**
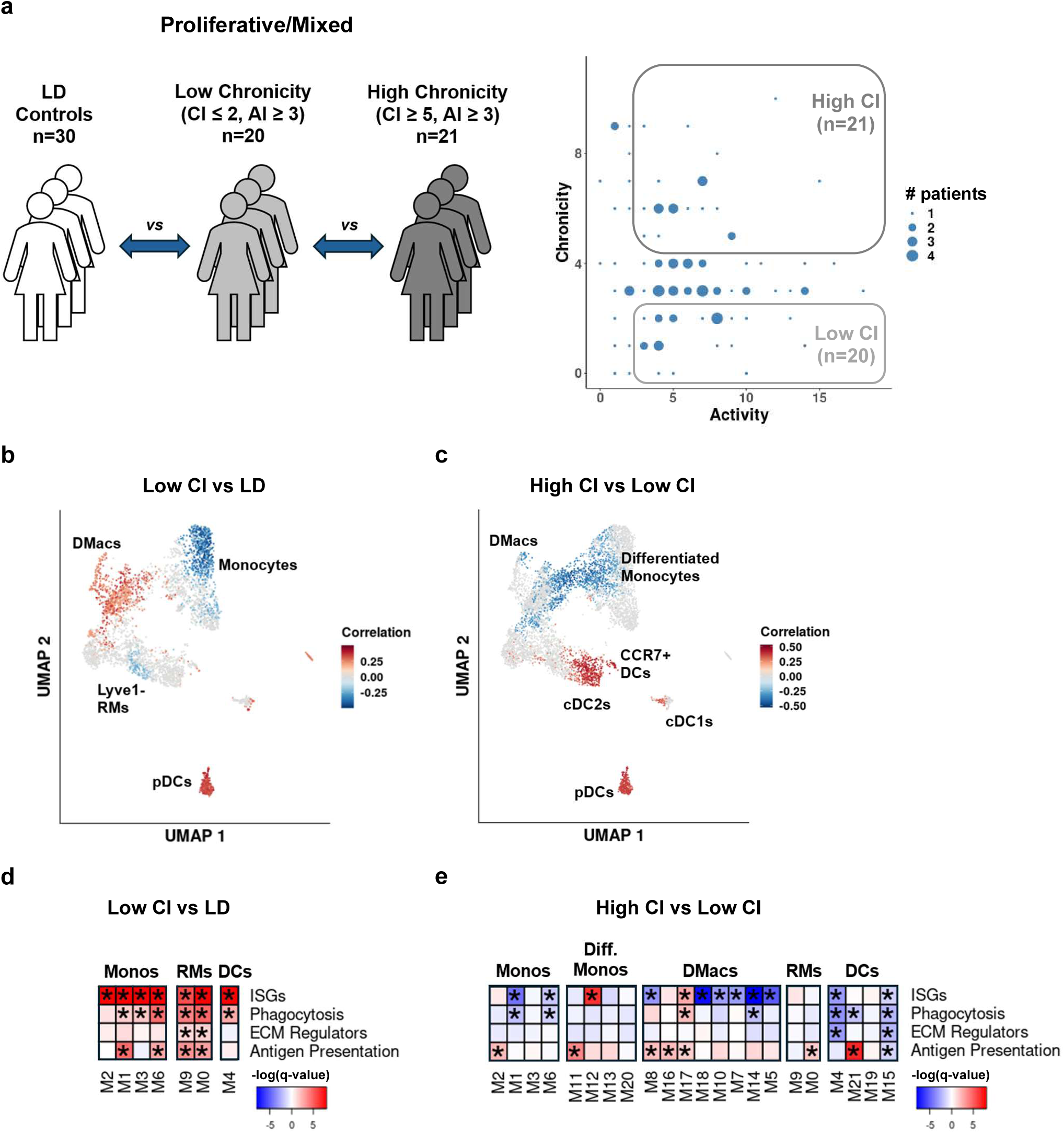
Analysis of proliferative/mixed LN. (a) The definition of the High CI and Low CI patient groups, for patients with proliferative/mixed LN (ISN class III, IV, III/V, IV/V). Each patient group is represented by a box, with the number of patients in the group specified in parentheses. Circle sizes correspond to the number of patients with specific values of AI and CI. (b-c) The results of running CNA; colored neighborhood had FDR ≤ 0.1. (b) Comparing the proliferative/mixed LN patients with Low CI (representing early disease) to LD controls. Cell populations expanded in LN patients are colored in red, those depleted in blue. (c) CNA results, comparing High CI (representing later stages of disease) and Low CI patient groups, for proliferative/mixed LN. Red represents expansion in the High CI group, compared to the Low CI patients; blue represents the opposite effect. (d-e) The results of running GSEA following DE analysis per cluster. Asterisks mark FDR ≤ 0.05. The color of each box was determined by the significance of the enrichment (darker shades mean more significant enrichment), and the sign of the enrichment, as specified below. (d) Comparison of Low CI proliferative/mixed LN patients and LD controls. Red designates higher expression in LN patients, blue – higher expression in LD controls. (e) Comparison of High CI and Low CI proliferative/mixed LN patients. Red – upregulated in the High CI group; blue – downregulated.

**Figure 4.**
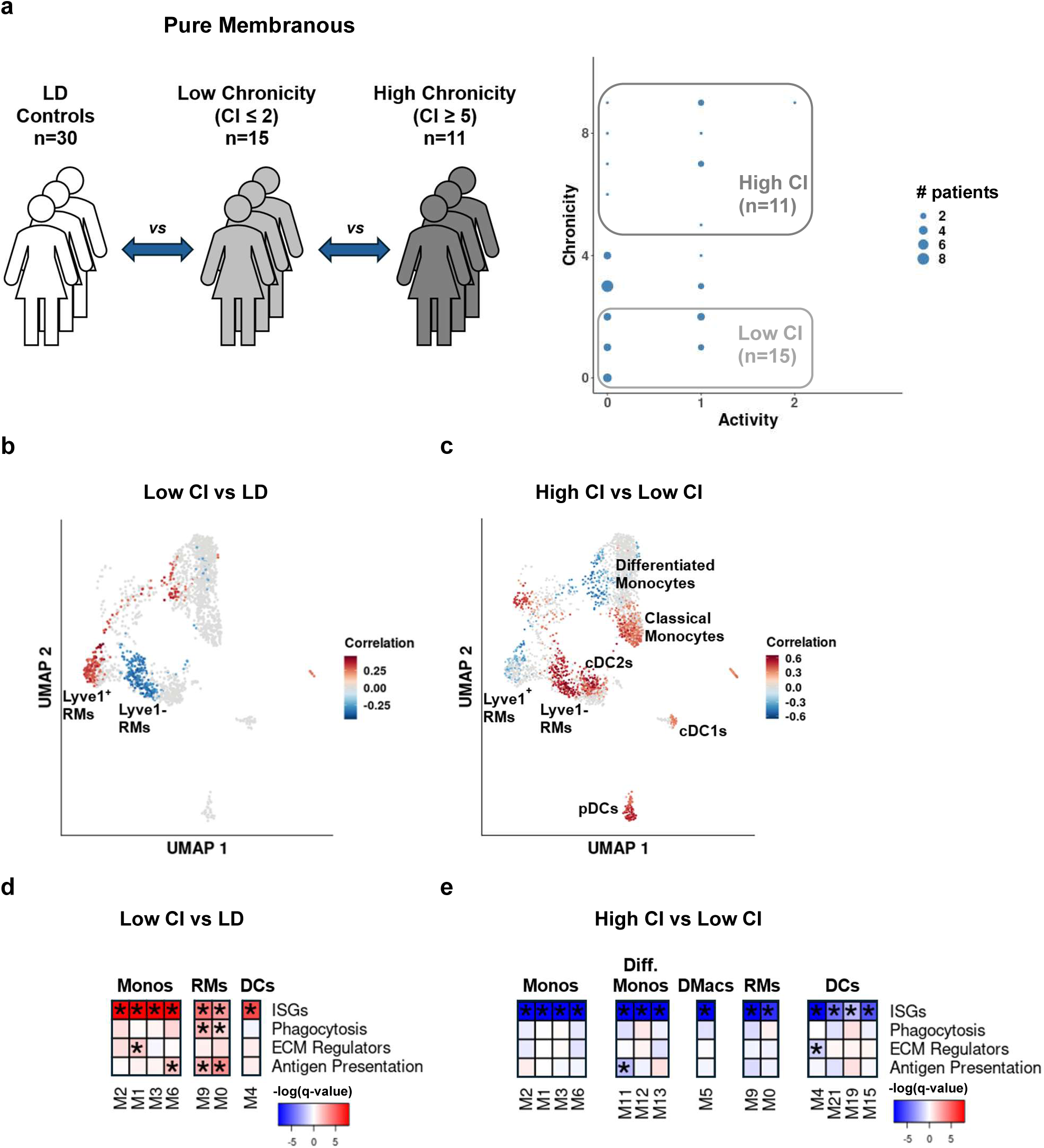
Analysis of pure membranous LN. (a) The definition of the High CI and Low CI patient groups, for patients with pure membranous LN (ISN class V). Each patient group is represented by a box, with the number of patients in the group specified in parentheses. Circle sizes correspond to the number of patients with specific values of AI and CI. (b-c) The results of running CNA; colored neighborhood had FDR ≤ 0.1. (b) Comparing the pure membranous LN patients with Low CI (representing early disease) to LD controls. Cell populations expanded in LN patients are colored in red, those depleted in blue. (c) CNA results, comparing the High CI (representing later stages of disease) and Low CI patient groups, for pure membranous LN. Red represents expansion in the High CI group, compared to the Low CI patients; blue represents the opposite effect. (d-e) The results of running GSEA following DE analysis per cluster. Asterisks mark FDR ≤ 0.05. The color of each box was determined by the significance of the enrichment (darker shades mean more significant enrichment), and the sign of the enrichment, as specified below. (d) Comparison of Low CI pure membranous patients and LD controls. Red – higher values in LN patients, blue – higher values in LD controls. (e) Comparison of High CI and Low CI pure membranous LN patients. Red represents upregulation in the High CI group, blue designates downregulation.

Co-varying neighborhood analysis (CNA) [20] showed that within the myeloid compartment, proliferative/mixed LN patients with active inflammation and low CI demonstrated a decrease in the relative frequencies of monocytes and an expansion of the DMac and differentiated monocyte clusters, compared to LD controls (Fig. 3b). Increase in the CI was associated with a decrease of the relative frequency of some of the DMac and differentiated monocyte clusters (M8, M10, M11 and M13) and an expansion of all DC clusters (Fig. 3c). Even after this relative contraction, the DMacs were still more abundant in the LN patients compared to LD controls (Supplementary Fig. 9a).

In pure membranous LN we observed only a minimal expansion of differentiated monocytes and DMacs in patients with low CI (Fig. 4b). As with proliferative/mixed LN, DC subsets expanded with the rise of the CI (Fig. 4c). A direct comparison of proliferative/mixed and pure membranous patients suggested that the latter patients did not differ much from LD controls in terms of their kidney myeloid cell frequencies, regardless of the CI (Supplementary Fig. 9b,c), and were characterized by relatively high levels of undifferentiated monocytes and low levels of DMacs.

### Low CI is associated with upregulated type I interferon response, phagocytosis and ECM regulation, while high CI involves a switch to antigen presentation

We next performed a differential expression (DE) analysis for each cluster, based on the patient groups defined above, followed by gene set enrichment analysis (GSEA) focusing on curated gene sets relevant for myeloid functions in LN (Supplementary Table 3). Compared to LD controls, patients with early disease (low CI) demonstrated an upregulation of ISGs and genes associated with antigen processing and presentation, in both proliferative/mixed and pure membranous disease (Figs. 3d, 4d). An increase in the CI was associated with a downregulation of the former genes and further upregulation of the latter ones, across most clusters (Figs. 3e, 4e). In the proliferative/mixed patients, early disease entailed an upregulation of genes associated with phagocytosis and ECM regulation, which later decrease. In early disease, the interferon response was higher in pure membranous compared to proliferative/mixed disease; once the CI increased this picture reversed, with ISGs returning to almost normal level in most clusters in membranous LN (Supplementary Fig. 10a-d). These changes in gene expression could only be partially explained by treatment with either mycophenolate mofetil (MMF) or high-dose corticosteroids (Supplementary Fig. 10e-h).

### Myeloid cells display preferential positioning within LN kidneys

To gain insights into the distribution of myeloid subsets within LN kidneys, we generated ST data from 6 LN patients and 2 healthy controls, utilizing a custom gene panel specifically designed to distinguish between the 24 myeloid subsets described in Fig 1a. To capture DMacs, we focused on patients with proliferative/mixed LN and low CI, in which our analysis suggests these subsets are expanded. As shown in Supplementary Fig. 11, the transcriptional signatures distinguishing between myeloid clusters in the scRNA-seq were effectively captured in the ST data.

We first observed that all myeloid subsets could be found in both the glomeruli and the tubulointerstitium (TI) (Fig. 5a). As expected, the density of immune cells in the TI parenchyma was significantly lower than in the inflamed parts of the TI (Supplementary Fig. 12a). We next checked if there are myeloid clusters demonstrating a statistically significant preference to occupy specific regions of the kidney (Fig. 5b; Supplementary Fig. 12b-h). Proinflammatory differentiated monocytes (clusters M12, M13) and some clusters of DMacs (M8, M16, M18) showed a preference for the glomeruli. Phagocytic DMac clusters (M7 and, to some extent, M10) and ISG-high monocytes/macrophages (M20) localized in periglomerular inflamed regions. In contrast, undifferentiated monocytes and RMs, including the related DMacs cluster M5, preferentially occupied the TI parenchyma (the latter observation held also in healthy controls). DCs concentrated in TI inflammatory aggregates.

**Figure 5.**
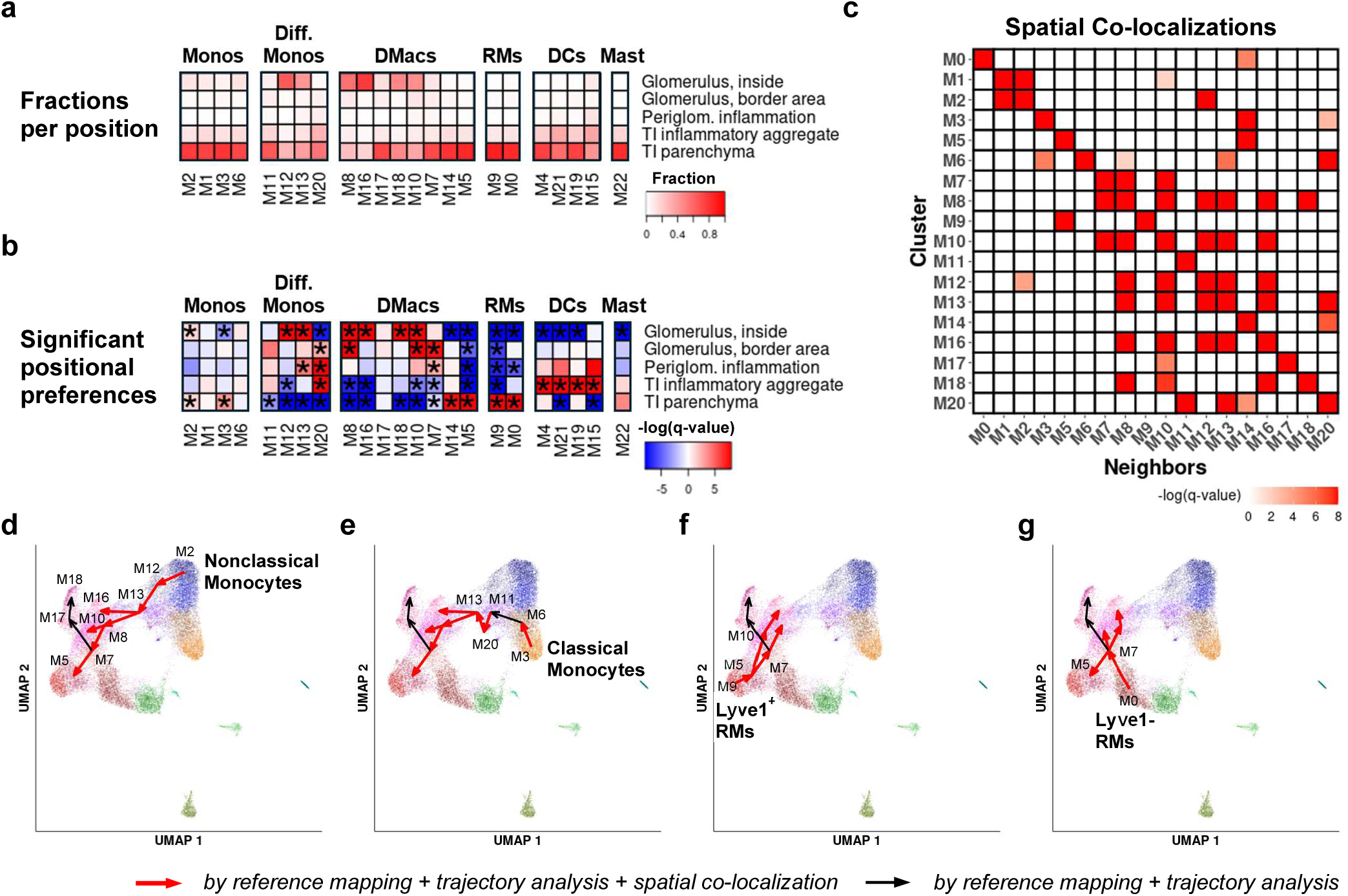
Spatial distribution and differentiation of myeloid clusters. (a) The spatial distribution of myeloid cells in kidneys of the 6 analyzed LN patients. Each row represents a specific region type (glomeruli, inflammatory aggregates in the TI etc.); each column represents a cluster, and shows the fraction of cells from that cluster in each region type (the sum of values per cluster is equal to 1). (b) The statistical significance of positional preferences, in the 6 analyzed LN kidneys. Each row represents a specific region type; each column represents a cluster. Heatmap elements are colored based on the magnitude of the adjusted p-values, and the direction of observed deviance from the null distribution assuming no preference: red represents a relative preference of a cluster to occupy a specific region type; blue represents a relative avoidance. Asterisks represent cases with FDR ≤ 0.05, further supported by the individual analysis of at least 2 separate study subjects. (c) Statistically significant co-localizations of cluster pairs in LN kidneys, with a focus on monocytes and macrophages. Each row pertains to a specific cluster, and shows co-localizations with other clusters of monocytes/macrophages. Colored heatmap elements represent cases with FDR ≤ 0.05, further supported by the individual analysis of at least 2 separate study subjects. Here, cells are considered adjacent if they are within 30μm of each other; similar results are found for choices of this distance threshold between 20μm and 50μm. (d-g) Assessing the feasibility of differentiation of monocytes and RMs into DMac clusters. Based on the results of trajectory analysis using PAGA, the shortest path from each putative origin into each DMac cluster is shown. Edges colored in red are also supported by the co-localization analysis presented in (c). (d) Paths originating from nonclassical monocytes (cluster M2). (e) Paths originating from classical monocytes (cluster M3). (f) Paths originating from Lyve1^+^ RMs (cluster M9). (g) Paths originating from Lyve1^−^ RMs (cluster M0). Note that the p-value of the edge between clusters M6 and M11 was just above the significance threshold. Edges connected to cluster M17 were not supported by the ST data, due to the small number of cells mapped to this cluster.

### Infiltrating monocytes and RMs likely differentiate into similar transcriptional states in LN kidneys

Comparison of LN myeloid cells with reference datasets suggested that DMac clusters comprise a mixture of cells derived from both infiltrating monocytes and RMs (Fig. 1d). This observation supports the hypothesis of “convergent differentiation”, in which both blood and tissue myeloid cells adopt similar transcriptional states as part of disease-associated changes in LN kidneys; a comparable phenomenon, in which Kupffer cells and infiltrating monocytes differentiate into similar disease-related macrophages, was recently described in the liver [21]. Trajectory analysis based on the scRNA-seq data supported this hypothesis, assigning high confidence scores to transitions between clusters allowing such convergent differentiation (Supplementary Fig. 13a, b).

To validate the predicted differentiation trajectories, we utilized our ST data, hypothesizing that if two clusters are related through differentiation, cells from both clusters will be spatially adjacent more often than expected by chance. The statistically significant co-localizations observed between cluster pairs (Fig. 5c; Supplementary Fig. 13c, d) supported most of the predicted transitions along the trajectories from the two monocyte states into each DMac cluster (Fig. 5d, e). Transitions from the two RM clusters, particularly from the Lyve1^−^ RMs (cluster M0) into the DMacs cluster M7, as well as the transitions from the DMac cluster M5 (which does co-localize with the Lyve1^+^ RMs) into the DMac clusters M7 and M10, were supported by the ST data when the analysis was confined to periglomerular inflamed regions. This indicates that these regions may serve as the preferred sites of differentiation from RMs (Fig. 5f, g; Supplementary Fig. 13e), whereas the differentiation from monocytes primarily occurs within the glomeruli.

To further explore the feasibility of monocyte differentiation into PECs, we incubated classical monocytes under various conditions and measured transcriptional changes using scRNA-seq. We found that stimulation by TGFβ induced the expression of genes associated with tissue injury, which are upregulated in most DMac clusters (Fig. 1e; Supplementary Fig. 13f). Stimulation by IFNγ had the opposite effect.

### Evidence for the modulation of PECs by DMacs

Previous studies have suggested that the activation, proliferation and migration of PECs is an essential step in replacing damaged podocytes and in the development of glomerulosclerosis [22, 23]. To explore the potential role played by DMacs in initiating these processes, we first redefined our patient groups, such that for the patients representing early disease, the glomerular components of the CI are equal to 0, i.e. before any irreversible glomerular damage is present; the results reported in Figs. 3 and 4 hold for this patient grouping as well (Supplementary Figs. 14, 15). We next performed DE analysis of PECs followed by GSEA, comparing proliferative/mixed LN patients with zero glomerular CI (gCI) to LD controls (Fig. 6a). Beyond the expected upregulation of ISGs, we identified a significant upregulation of pathways previously demonstrated to be associated with a migratory phenotype in epithelial cells [24]. Notably, these changes were not observed in pure membranous patients (Supplementary Fig. 16a), in which there is less glomerular inflammation and little DMac expansion. Out of the genes included in these pathways, 58 were significantly upregulated in our data (Supplementary Table 3); we use these genes to define a “migration score”, employed in subsequent analyses.

**Figure 6.**
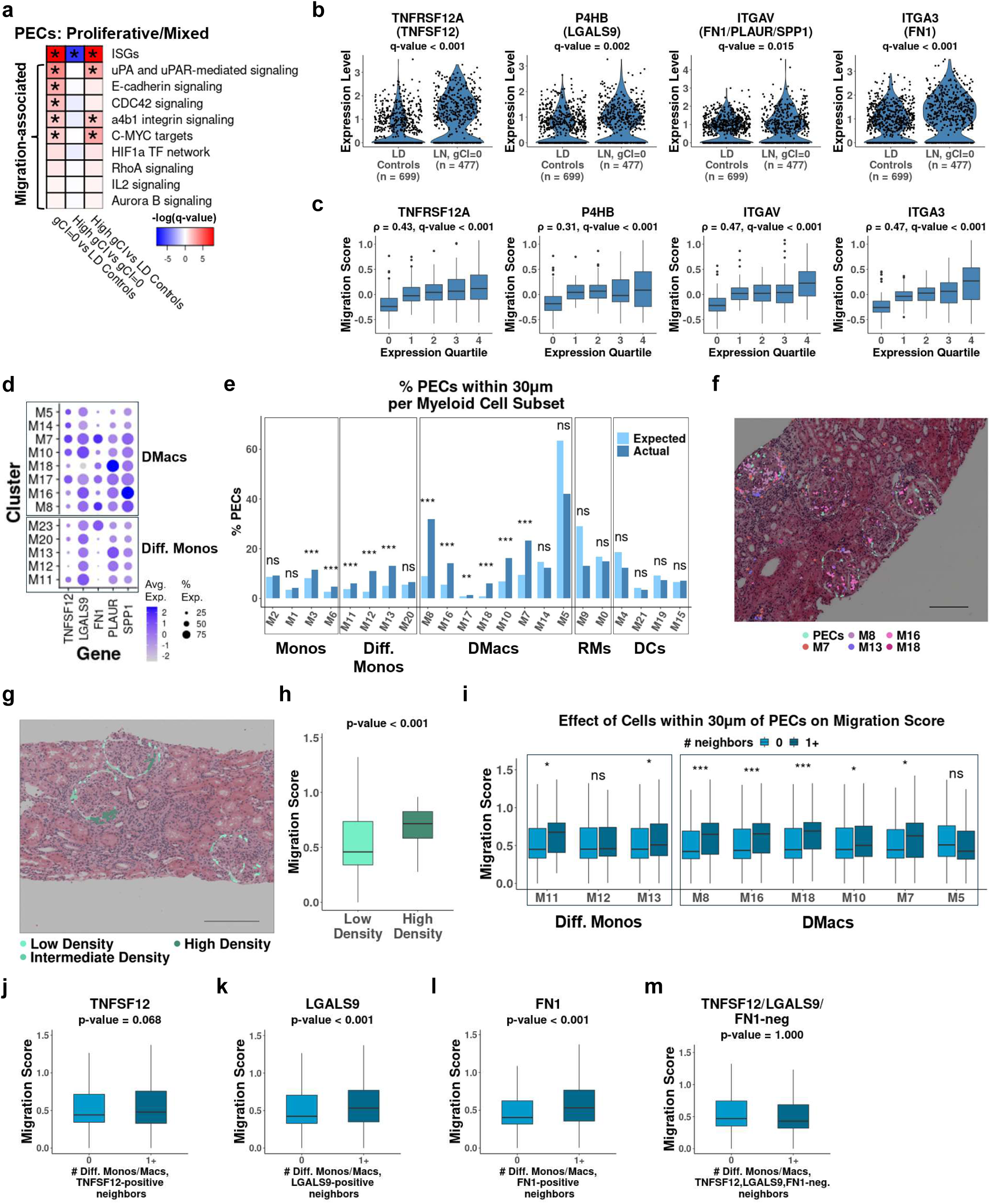
Putative interactions of PECs and myeloid cell subsets. (a) The results of running GSEA following DE analysis for PECs, comparing: (i) the gCI=0 proliferative/mixed patients and LD controls (left; red means upregulation in LN patients); (ii) High gCI and gCI=0 proliferative/mixed patients (middle; red denotes upregulation in the High gCI patients), and (iii) High gCI proliferative/mixed patients and LD controls (right; red represents upregulation in LN patients). The glomerular CI (gCI) is computed as the sum of components of the NIH chronicity index that pertain to the glomeruli, i.e. global or segmental glomerulosclerosis, and fibrous crescents. Tested were ISGs as well as gene sets previously shown to be associated with epithelial cell migration. Asterisks mark FDR ≤ 0.05. The color of each box was determined by the significance of the enrichment (darker shades mean more significant enrichment) and the sign of the enrichment. (b) Changes in the expression levels in PECs of several receptors, comparing the gCI=0 proliferative/mixed patients and LD controls. For each gene, the FDR-adjusted p-value (q-value) is specified. n denotes the number of cells included in each group. The corresponding ligands are specified in parentheses. (c) The relation between the expression level of each receptor shown in (b) and the migration score, in each PEC. Included are only PECs from gCI=0 proliferative/mixed LN patients. Cells were grouped into bins based on the expression level of the receptor shown in each panel; one bin includes cells with an expression value of 0, while the rest of the bins correspond to the quartiles of the expression distribution of the receptor, taking into account only the positive expression values. For each panel, the Spearman correlation ρ and its associated p-value are reported, taking into account all PECs from gCI=0 proliferative/mixed LN patients. (d) The expression, in differentiated monocyte and DMac clusters, of ligands binding the receptors appearing in (b). (e) The percentage of PECs adjacent to each subset of myeloid cells, within a distance of 30μm, in the spatial transcriptomics data. Dark blue shows the actual percentage, light blue the percentage expected if immune cells were distributed uniformly in the tissue. *** - FDR < 0.001; ** - FDR < 0.01; ns – not significant. (f) A representative example of co-localization of PECs and selected subsets of myeloid cells, in a slide taken from the spatial transcriptomics data. Cell positions are shown overlaid on H&E staining. Scale bar denotes 200μm. (g) A representative example of PEC aggregates, in a slide taken from the spatial transcriptomics dataset. Shown are PECs, colored by the number of PECs in their immediate neighborhoods (with a threshold set arbitrarily to 30μm), and overlaid with . Light green – PECs from low density neighborhoods (less than 5 adjacent PECs); green – intermediate density neighborhoods (5 or more, less than 8); dark green – high density neighborhoods (PEC aggregates; 8 or more adjacent PECs). The reported results were found to be robust with regard to the actual choice of these thresholds. Scale bar denotes 200μm. (h) The migration score, in PECs found in low-density and high-density environments. The p-value was computed using the Mann-Whitney U test. (i) The effect of neighboring cells from selected myeloid clusters on the migration score in PECs. Here, cells are considered adjacent if they are within 30μm of each other. For each myeloid cluster, light blue designates the case of PECs with no neighbors of that cluster, and dark blue represents the case where there is at least one such neighbor. P-values were computed using the Mann-Whitney U test, then corrected for multiple comparisons using the Benjamini-Hochberg method. *** - FDR < 0.001; * - FDR < 0.05; ns – not significant. (j-m) The change in the migration score in PECs, based on the absence or presence of adjacent DMacs or differentiated monocytes expressing a given gene. P-values were calculated using a unidirectional Mann-Whiteny U test, checking an increase in the migration score. (j) The effect of adjacent DMacs or differentiated monocytes expressing TNFSF12 (TWEAK). (k) The effect of adjacent DMacs or differentiated monocytes expressing LGALS9 (Galectin 9). (l) The effect of adjacent DMacs or differentiated monocytes expressing FN1 (Fibronectin 1). (m) The effect of adjacent DMacs or differentiated monocytes not expressing TNFSF12, LGALS9 and FN1.

Next, we looked for differentially expressed receptors in PECs in the above analysis, such that: (i) the ligand binding these receptors is expressed in DMac or differentiated monocyte clusters that are enriched in or around the glomeruli, and (ii) the binding of the receptor was previously shown to be associated with the activation, proliferation and/or migration of epithelial or endothelial cells. We found several such receptors and ligands (Table 1; Fig. 6b,d, Supplementary Fig. 16b,d). Interestingly, for all these receptors we observed a statistically significant positive correlation between their expression in PECs and the expression of the migration score in the same cells (Fig. 6c, Supplementary Fig. 16c), suggesting that signaling through these receptors may be part of the events leading to PEC migration. Of note, these receptors were not differentially expressed in pure membranous disease (Supplementary Fig. 16e).

**Table 1.** Receptor-ligand pairs potentially mediating interactions between PECs and differentiated monocytes/DMacs.

| **Receptor in**<br>**PECs** | **Ligands in Diff.**<br>**Monocytes and**<br>**DMacs** | **Functional Effect** | **References** |
| --- | --- | --- | --- |
| TNFRSF12A | TWEAK<br>(TNFSF12) | Activation, proliferation,<br>migration | [25-28] |
| P4HB | Galectin 9<br>(LGALS9) | Proliferation, migration | [29, 30] |
| ITGAV | Fibronectin 1<br>(FN1), PLAUR,<br>Osteopontin<br>(SPP1) | Activation, proliferation,<br>migration | [31-33], [34-36], [37-40] |
| ITGA3,<br>ITGB5 | Fibronectin 1 (FN1) | Activation, proliferation,<br>migration | [31-33] |
| ITGB1 | Fibronectin 1<br>(FN1), COL8A2 | Activation, proliferation,<br>migration | [31-33], [41] |
| AXL | GAS6 | Proliferation, migration | [42-44] |
| DPP4 | CXCL2, CXCL10 | Modulation of activation<br>and proliferation | [45-47] |

These results imply a possible role for DMacs and differentiated monocytes in promoting PEC activation, proliferation and migration. We utilized our ST data to further validate this prediction. We first observed that a substantial fraction of PECs was found in the vicinity of several clusters of DMacs and differentiated monocytes; furthermore, this occurred more than expected by chance, i.e. if immune cells were spread uniformly across the tissue (Fig. 6e,f; Supplementary Fig. 17a). This observation held over a wide range of distance thresholds defining cellular adjacency (Supplementary Fig. 17b-c).

Next, we found that PECs may form aggregates (i.e. areas of increased cell density; Fig. 6g; Supplementary Fig. 18a). PECs present in such aggregates had a significantly elevated migration score (Fig. 6h), indicating that this metric indeed reflects histologic abnormalities. In addition, such PECs had a significantly lower interferon score (Supplementary Fig. 18b), in accordance with the observation from the scRNA-seq data showing that the formation of irreversible tissue damage is associated with a decrease in the expression of ISGs in PECs (Fig. 6a).

To test whether DMacs and differentiated monocytes may induce the formation of PEC aggregates, we focused on the neighborhoods of PECs outside of aggregates (i.e. prior to aggregate formation), and checked the effect on the migration score of having a neighbor from each relevant myeloid subset. Several clusters were associated with a significant increase in the migration score, including M7, M8, M13, M16 and M18, regardless of the distance threshold used to define cellular adjacency (Fig. 6i; Supplementary Fig. 18c,d). Testing directly the potential role of specific ligands in the induction of PEC aggregates, we identified a statistically-significant increase of the migration score in PECs adjacent to DMacs and differentiated monocytes expressing TNFSF12 (TWEAK), LGALS9 (Galectin 9) or FN1 (Fibronectin 1), as compared to DMacs/differentiated monocytes lacking these 3 ligands (Fig. 6j-m).

## DISCUSSION

In this work, we have generated a large-scale scRNA-seq dataset of LN kidney samples, allowing us to construct a high-resolution comprehensive catalog of all myeloid cell subsets and activation states found in patients, representing a range of histologic classes and LN-associated tissue injury. Utilizing the intrinsic heterogeneity of our cohort and detailed clinical annotations coupled to it, together with comparisons to published reference datasets, a focused application of spatial profiling of LN kidneys and the performance of in vitro culturing experiments, we were able to derive detailed hypotheses linking immunologic mechanisms to histopathologic features of LN.

Our analyses suggest that the early stage of LN, characterized by low CI, is marked by increased frequencies of monocytes and macrophages and exhibit a strong type I interferon response. In proliferative/mixed LN, this stage is also associated with DMac differentiation and upregulation of genes involved in phagocytosis. We propose that as the CI rises, the immune response in LN kidneys demonstrates a gradual increase in the presence of B cells, CD4^+^ memory T cells and DCs, along with a decreased type I interferon response. This switch is accompanied by upregulation of genes related to antigen processing and presentation in macrophages and DCs. Furthermore, we recently showed that the increase in CI is in addition associated with a decrease in CD56dim NK cells and CTLs, and a parallel increase in stem-like GZMK^+^CD8^+^ T cells (Al Souz et al; manuscript under review). These observations suggest that as irreversible tissue damage accumulates in LN kidneys, the local immune response gradually shifts from one dominated by monocytes, macrophages and cytotoxic cells, centered mainly in the glomeruli, to one featuring increased antibody production modulated by T helper cells and supported by DCs, taking place in the TI.

We designed the gene panel used to generate our ST data such that it would capture all myeloid subsets found in the scRNA-seq dataset. Compared to recent publications using ST to profile LN kidneys [48–50], this allowed us to increase the level of detail of our analysis. In particular, we could identify preferential positional patterns of specific myeloid cell subsets; use the ST data to validate predicted differentiation trajectories; and point to specific myeloid subsets likely interacting with PECs.

There are a number of limitations to our study. First, the processing of kidney samples for scRNA-seq may skew the estimated cluster frequencies, as not all cells survive freezing, thawing and tissue dissociation equally. Second, our dataset includes only a single time point per patient. Nonetheless, we could utilize the size and heterogeneity of our cohort, and the fact that irreversible tissue damage generally accumulates over time, to point to possible cellular and molecular events associated with disease progression. Indeed, as we have shown above, the ratio between the relevant lymphoid and myeloid cells was correlated with patients’ age at the time of biopsy (Supplementary Fig. 2f-g). A definitive description of disease processes would require serial biopsies, which is beyond the scope of the current study; still, our observations may serve as hypotheses to longitudinal follow-up studies.

We could not identify effective cellular and molecular predictors of treatment response in the kidneys of LN patients, beyond those reflecting the CI; similarly, given the CI, we did not observe significant differences between biopsies acquired at first onset of LN and those obtained at disease relapse (Supplementary Fig. 19a-d). One possible explanation of this is the relatively strong signal correlated with the CI, which masks weaker signals. In addition, the fact that treatment was not uniform across the studied cohort likely increases the noise in our data. Furthermore, multiple factors may contribute to non-response downstream of the biopsy including access to medical care, hypertension and drug adherence. As the present study was not a clinical trial, these factors were not controlled.

Our work significantly enriches the knowledge of disease mechanisms in human LN. As such, it can benefit future efforts to develop novel therapies for this disease, while providing a detailed dataset allowing to test new hypotheses regarding the cell populations and molecular pathways driving it.

## Supporting information

Supplementary Table 1 - Cohort

Supplementary Table 2 - Myeloid Cluster Markers

Supplementary Table 3 - Gene Lists

Supplementary Table 4 - Clusters Ratio vs CI

Supplementary Table 5 - DASH primer sequences

Supplementary Table 6 - Cluster Merging Schemes

## ACKNOWLEDGEMENTS AND AFFILIATIONS

We thank the patients who participated in this study, and the scientists and clinical sites in the Accelerating Medicines Partnerships in RA/ SLE. This work was supported by the Accelerating Medicines Partnership® Rheumatoid Arthritis and Systemic Lupus Erythematosus (AMP® RA/SLE) Network (see below). AMP is a public-private partnership (AbbVie, Arthritis Foundation, Bristol-Myers Squibb Company, Foundation for the National Institutes of Health, GlaxoSmithKline, Janssen Research and Development, LLC, Lupus Foundation of America, Lupus Research Alliance, Merck & Co., Inc., National Institute of Allergy and Infectious Diseases, National Institute of Arthritis and Musculoskeletal and Skin Diseases, Pfizer, Inc., Rheumatology Research Foundation, Sanofi and Takeda Pharmaceuticals International, Inc.) created to develop new ways of identifying and validating promising biological targets for diagnostics and drug development. Accelerating Medicines Partnership and AMP are registered service marks of the US Department of Health and Human Services. Funding was provided through grants from the National Institutes of Health: UH2-AR067676 (AMP RA/SLE), UH2-AR067677 (AMP RA/SLE), UH2-AR067679 (AMP RA/SLE), UH2-AR067681 (AMP RA/SLE), UH2-AR067685 (AMP RA/SLE), UH2-AR067688 (AMP RA/SLE), UH2-AR067689 (AMP RA/SLE), UH2-AR067690 (AMP RA/SLE), UH2-AR067691 (AMP RA/SLE), UH2-AR067694 (AMP RA/SLE), UM2-AR067678 (AMP RA/SLE), RO1-DK131482-01.

## AMP RA/SLE Network

### Operation/Scientific

Operations – William Apruzzese, Jennifer Goff, Patrick J. Dunn

SBG – Soumya Raychaudhuri, Fan Zhang, Ilya Korsunsky, Aparna Nathan, Joseph Mears, Kazuyoshi Ishigaki, Qian Xiao, Nghia Millard, Kathryn Weinand, Saori Sakaue

LC/STAMP – PJ Utz, Rong Mao, Bill Robinson, Holden Maecker

OMRF – Judith James, Joel Guthridge, Wade DeJager, Susan Macwana

### SLE

Rochester – Jennifer Anolik, Jennifer Barnas

BWH – Michael B. Brenner, James Lederer, Deepak A. Rao

Northshore – Betty Diamond, Anne Davidson, Arnon Arazi

UCSF – David Wofsy, Maria Dall’Era, Raymond Hsu

NYU – Jill Buyon, Michael Belmont, Peter Izmirly, Robert Clancy, Phillip Carlucci, Kristina Deonaraine

Einstein – Chaim Putterman, Beatrice Goilav

Hopkins – Michelle Petri, Andrea Fava, Jessica Li

MUSC – Diane L. Kamen

Cincinnati – David Hildeman, Steve Woodle

Broad – Nir Hacohen, Paul Hoover, Thomas Eisenhaure, Michael Peters, Arnon Arazi, Tony

Jones, David Lieb

Rockefeller – Thomas Tuschl, Hemant Suryawanshi, Pavel Morozov, Manjunath Kustagi

UCLA – Maureen McMahon, Jennifer Grossman

UCSD – Ken Kalunian

Michigan – Matthias Kretzler, Celine Berthier, Jeffery Hodgin, Raji Menon

Texas Tech – Fernanda Payan-Schober, Sean Connery

Cedars – Mariko Ishimori, Michael Weisman

## AUTHORSHIP CONTRIBUTION STATEMENT

**Paul J. Hoover:** Formal analysis, Conceptualization, Methodology, Investigation, Data curation, Writing – original draft, Writing – review & editing, Visualization. **Thomas M. Eisenhaure:** Methodology, Investigation, Writing – original draft. **Jeffrey Hodgin:** Data curation. **William Apruzzese:** Data curation. **Joseph Mears:** Formal analysis. **Michael Peters:** Investigation. **Tony Jones:** Investigation. **Sujal I. Shah:** Data curation. **Haniya Kamal:** Formal analysis. **Rollin Leavitt:** Investigation. **Shaun W. Jackson**: Conceptualization, Resources. **Patrick Danaher:** Conceptualization, Resources. **Deepak A. Rao:** Writing – review & editing. **Qian Xiao:** Formal analysis. **Siddarth Gurajala:** Formal analysis. **Andrea Fava:** Resources, Writing – review & editing. **Celine C. Berthier:** Writing – review & editing. **Alice Horisberger:** Writing – review & editing. **Peter M. Izmirly:** Resources, Data curation, Writing – review & editing. **H. Michael Belmont:** Resources, Data curation. **Robert Clancy:** Resources, Data curation. **Richard Furie:** Resources. **Joel M. Guthridge:** Resources. **Maria Dall’Era:** Resources. **David Wofsy:** Funding acquisition, Resources. **Diane L. Kamen:** Resources. **Kenneth C. Kalunian:** Resources. **Maureen A. McMahon:** Resources. **Jennifer Grossman:** Resources. **Fernanda Payan-Schober:** Resources. **David A. Hildeman:** Resources. **E. Steve Woodle:** Resources. **Chaim Putterman:** Funding acquisition, Resources. **Matthias Kretzler:** Conceptualization, Resources. **Soumya Raychaudhuri:** Conceptualization, Supervision. **Judith A. James:** Funding acquisition, Resources, Writing – review & editing. **Jennifer H. Anolik:** Funding acquisition, Resources, Writing – review & editing. **Michelle A. Petri:** Funding acquisition, Resources, Writing – review & editing. **Jill P. Buyon:** Funding acquisition, Resources, Writing – review & editing. **The Accelerating Medicines Partnership (AMP) RA/SLE Network**: Resources. **Betty Diamond:** Supervision, Conceptualization, Funding acquisition, Writing – review & editing, Resources. **Anne Davidson:** Supervision, Conceptualization, Funding acquisition, Writing – review & editing, Resources. **Nir Hacohen:** Supervision, Conceptualization, Funding acquisition, Writing – review & editing. **Arnon Arazi:** Supervision, Conceptualization, Formal analysis, Writing – original draft, Writing – review & editing, Visualization.

## Supplementary Information

### EXTENDED METHODS

#### Patient recruitment

Details of enrollment into the AMP SLE cohort have been previously published [1–3]. In brief, patients with LN undergoing kidney biopsies as part of standard of care were eligible to enroll in the prospective AMP LN study. The decision to biopsy was at the discretion of the treating physician to confirm suspected LN de novo or relapse of disease or activity not responding to treatment. Inclusion in AMP required: 1) age ≥ 18; 2) fulfilment of the revised American College of Rheumatology [4, 5] or the Systemic Lupus Erythematosus International Cooperating Clinics [6] classification criteria for SLE; 3) a urine protein/creatinine ratio (UPCR) >0.5 at the time of biopsy. For the analyses reported herein, classification of responder status was restricted to patients with baseline random or 24-hour UPCR ≥ 1.0 since for patients with ratios between 0.5 and 0.999, proteinuric response has not been defined. Only patients with renal biopsies that demonstrated International Society of Nephrology/Renal Pathology Society (ISN/RPS) classes III, IV, V or combined III or IV with V read by the pathologist at each participating site were considered in this analysis. At a later stage, however, biopsies were scored by a central pathologist (JH), to achieve uniformity in downstream analyses. Central scoring was used when available; otherwise, the scoring from the participating site was used. Exclusion criteria included: 1) a history of kidney transplant; 2) rituximab treatment within 6 months of biopsy; 3) pregnancy at the time of biopsy. The study protocol was approved by the institutional review boards and ethics committees of participating sites in adherence with the Declaration of Helsinki and all informed consent was obtained from all participants.

#### Clinical metadata

Clinical metadata from all patients was recorded at the time of biopsy, from a predetermined set of categories, including self-reported race (Asian, Black, White, Other)/ethnicity (Hispanic, non-Hispanic) as required for NIH-funded studies. Medications and laboratory tests were documented at each visit (baseline, week 12, week 26 and week 52) and were performed at the participating sites. Given that medication changes occurred after the baseline visit in response to the kidney biopsy results, we chose the week 12 treatment to represent the induction regimen and regarding steroids, the higher dose at either baseline or week 12 was considered the induction dose for similar reasons. Pulse steroids were also captured separately.

Treatment outcome was determined as follows. For complete response (CR) we required: 1) UPCR < 0.5 (variations on these threshold, set to either 0.7 or 0.8, were also tested in downstream analyses); and 2) normal creatinine (≤ 1.3 mg/dL) or, if abnormal, ≤ 125% of baseline; and 3) prednisone ≤ 10 mg/day at the time of the study visit. Partial response required: 1) > 50% reduction in UPCR; and 2) normal creatinine (≤ 1.3 mg/dL) or, if abnormal, ≤ 125% of baseline; and 3) prednisone dose ≤ 15 mg/day at the time of the study visit. Patients who did not achieve a CR or PR at the specific timepoints were considered non-responders (NR) or not determined (ND) if data were missing. These response definitions were based on the ACCESS Trial [7]. In agreement with the ACCESS trial, we specifically decided not to include the microscopic review of the urine sediment given the absence of uniformity across sites in assessing urinary sediment and the challenge of attribution especially in a population of menstruating women. The prednisone threshold for CR at ≤ 10mg prednisone was also based on the ACCESS trial. However, the ≤ 15 mg prednisone maximum for defining PR was agreed upon unanimously by the site investigators. Although proteinuria was measured by either a UPCR on a spot urine or a timed urine collection, consistency of the method across the study for an individual was required. While determination from a timed urine collection was preferred, if this method was not performed at all time points for an individual participant, calculations from a spot urine were utilized.

#### Kidney single-cell RNA sequencing

Human kidney needle biopsies were cryopreserved in Cryostor CS10 (Stemcell Technologies) at sites of clinical care, and shipped to Oklahoma Medical Research Foundation Biorepository for storage in liquid nitrogen. Samples were randomized and shipped on dry ice to the Broad Institute for dissociation and sequencing.

Biopsies were processed in batches of 3. Samples were thawed at 37C and then transferred to RPMI (ThermoFisher) supplemented with 10% heat-inactivated FCS (Avantor) and 10mM HEPES (ThermoFisher) for 1 minute at room temperature. Next, the samples were placed in 100uL RPMI with 0.04% BSA (Millipore Sigma) and 10mM HEPES and cut into ∼2mm segments (From some biopsies that were at least 7mm in length two segments were used to isolate nuclei and perform single nucleus RNA-seq [8]. These data are not used in this study since the number of immune cells in single nucleus RNA-seq data was relatively low). The tissue segments and RPMI/HEPES/BSA were then added to 500uL DMEM/F12 media (Corning) with 0.5mg/mL Liberase TL (Roche) and agitated in an orbital mixer set to 300RPM and 37 degrees C for 12 minutes. Midway through digestion the samples were mixed by gentle pipetting with a large-bore tip (Rainin). Following digestion the samples were placed on ice, and 600uL cold RPMI/FCS/HEPES was added. The sample was then strained through a 70um mesh (Miltenyi). The strainer was rinsed with 4mL cold RPMI/BSA/HEPES. Remaining tissue fragments on the strainer were mechanically homogenized using a cell strainer pestle (CELLTREAT), following which the strainer was rinsed with an additional 5mL cold RPMI/BSA/HEPES. All strained sample was collected in a conical tube and pelleted at 200 x g for 8 minutes at 4 C. Pellets were resuspended in ∼50uL RPMI/BSA/HEPES and strained through 40um mesh (Pluriselect). The resulting cell suspension was transferred to a clean tube and counted by trypan blue exclusion. Up to 10,000 live cells per sample were loaded in separate 10x channels and profiled using Chromium 3’ V3 single cell gene expression kits (10x Genomics). Sequencing libraries were generated by the Broad Genomics Platform according to manufacturer’s specifications.

To reduce sequencing requirements, we used the DASH method [9], with modifications for use with 3’ 10x libraries. A panel of 61 guide RNA sequences (Supplementary Table 5) tiling mitochondrial transcript regions found in the kidney RNA-seq libraries was designed using the Broad Institute Genetic Perturbation Platform algorithm [10, 11] to predict on-target and off-target activity. Two mitochondrial transcripts, *MT-ND5* and *MT-ND6,* with relatively low expression were exempted from targeting to allow their expression to be used as a proxy for overall mitochondrial content during QC and analysis. Oligonucleotides (IDT) containing the guide RNA sequences (Supplementary Table 5) were pooled and used to generate template for reverse transcription as previously described [9]. Guide RNA was then transcribed by incubation of 75ng template DNA for 4 hours at 37C with 100U T7 RNA polymerase (New England Biolabs) in a 30uL reaction containing 1X RNAPol Reaction Buffer and 0.83 mM each of rATP, rCTP, rGTP and rUTP (New England Biolabs). After removing template DNA by digestion with 4U DNAseI (New England Biolabs) for 15 minutes at 37C, RNA was purified using a Monarch RNA Cleanup Kit (New England Biolabs) and quantified by absorbance of 260nm light. Cas9:gRNA complexes were produced by mixing 1.5 pmoles Cas9 nuclease (New England Biolabs) with 1.5 pmoles of the gRNA pool in a 15 uL solution containing 2uL 10 X NEBuffer™ r3.1 (NEB) and incubating at 25C for 15 minutes. Targeted double strand breaks were then made in gene expression sequencing libraries by adding 10ng of a single library in 5uL water to the ribonucleoprotein (RNP) complexes and incubating for 2h at 37C. RNPs were removed from the digested libraries to enable direct amplification [12]; the samples were incubated with 1ug RNAseA (Ambion) and 0.8U Proteinase K (NEB) for 15 minutes at 37C, then for 15 minutes at 95C to inactivate the enzymes. The libraries were then amplified by 6 PCR cycles using P5/P7 oligonucleotides (AATGATACGGCGACCACCGA, CAAGCAGAAGACGGCATACGA) and KAPA HiFi HotStart ReadyMix (Roche). Finally, the PCR products were purified using a 0.8x volumetric ratio of SPRIselect paramagnetic beads (Beckman Coulter). The resulting libraries were quantified and sequenced on a Novaseq S4 (Illumina).

#### scRNA-seq analysis

Following alignment, basic scRNA-seq processing, including QC and clustering, was performed using Seurat v4 [13]. Cells with less than 500 detected genes, less than 1000 UMIs or a percentage of reads mapped to mitochondrial genes (following DASH) larger than 3% were discarded. Batch correction was done using Harmony [14], taking into account the sample collection site, the processing batch and the individual sample itself. Graph-based clustering was performed in a step-wise manner, with increasing resolution: first, we identified clusters of immune and non-immune cells. Immune cells were then clustered separately, in order to identify general lineages – myeloid cells, T/NK cells, B cells and plasma cells. Dividing cells were also identified as a separate population. Each lineage was then clustered separately, to identify high-resolution cell subsets. In each case, the clustering resolution was chosen based on the expression patterns of known canonical markers, such that groups of cells on the UMAP plane demonstrating clearly distinct expression patterns would be included in separate clusters. Furthermore, we took into account mapping to reference datasets, aiming for clusters demonstrating relatively uniform mapping results when possible. For non-immune cells, we took a more focused approach, as our goal here was to identify PECs only.

Cluster labeling was based on the expression of canonical markers, reference mapping to published scRNA-seq datasets, and the genes distinguishing each cluster when compared to the others. Reference mapping was performed using Symphony [15]. The mapping results were validated using the reference mapping methods provided by Seurat, as well as the Pearson correlation-based approach previously presented [16]; all 3 methods provided similar results, confirming the validity of the mapping. For the mapping to the reference dataset taken from Villani et al. [17], we report the mapping results using the Pearson correlation-based approach, as in our hands it performs better for small reference datasets. When mapping to the reference dataset from Stewart et al. [18], we changed the original reference label ‘DC’ to ‘RM1/DC’, as the comparison to blood cells suggested that this reference class is a mixture of RMs and DCs. To distinguish between DC subsets, all clusters in which a large percentage of cells were mapped to the DC category in Stweart et al. [18]were further mapped to the blood reference dataset from Villani et al. [17].

#### Computation of composite gene expression scores

To compute composite scores representing the expression levels of pre-defined gene sets, we calculated the average of the scaled expression across all genes contained in the set. The genes included in each set are provided in Supplementary Table 3. The same approach was used for both the scRNA-seq and ST data.

The migration score was based on a previous publication [19], considering all pathways reported there. We then included only pathways enriched in GSEA when comparing proliferative/mixed LN patients with gCI=0 to LD controls. From these pathways, we further included only genes found to be actually differentially expressed in that comparison. The final list of genes is found in Supplementary Table 3.

#### Principal component analysis (PCA) of cluster frequencies

PCA was performed based on the fraction of each cluster out of all immune cells, without scaling. Before PCA, clusters were merged to consolidate small clusters. We further included only patients with at least 20 immune cells in their kidneys. For sensitivity analysis, we tested several thresholds on immune cell numbers in the range 10-100 cells; furthermore, several cluster merging schemes were considered (Supplementary Table 6); in all cases, the general results reported were conserved.

#### Computing cluster frequency ratios

For the ratio between memory CD4^+^ T cells, GZMK^+^CD8^+^ T cells and B cells and between monocytes and macrophages, the latter group included clusters M1, M2, M3, M5, M6, M7, M8, M10, M11, M12, M13, M14, M16, M17, M18, M20 and M23. When computing the ratio between DMacs and monocytes, the numerator included clusters M5, M7, M8, M10, M14, M16, M17 and M18, and the denominator clusters M1, M2, M3 and M6. In all ratio calculations, we included only samples which had at least 20 cells overall for both the numerator and denominator; the reported results were robust for a wide range of choices for this threshold.

#### Definition of patient groups

Patient groups were defined separately for proliferative/mixed LN and pure membranous LN based on the NIH chronicity index (CI). For patients with proliferative/mixed LN, additional threshold with regard to the NIH activity index (AI) were imposed. Several choices for these thresholds were tested to check the robustness of the results, as reported in the text. When running DE analysis for PECs (Fig. 6, Supplementary Fig. 14, 15), we defined the patient groups based on the glomerular components of the NIH chronicity index (global or segmental glomerulosclerosis, fibrous crescents) and the glomerular components of the NIH activity index (endocapillary hypercellularity, neutrophils karyorrhexis, wire loops or hyaline thrombi, fibrinoid necrosis, cellular or fibrocellular crescents).

#### Co-varying neighborhood analysis (CNA)

CNA [20] was run using the R implementation. Results with FDR ≤ 0.1 were reported.

#### Differential expression (DE) analysis and gene set enrichment analysis (GSEA)

DE analysis was performed using a generalized linear mixed-effects model (GLMM) using the R package glmer, following a previously described approach [21]. Gene expression in each cell was modeled using a negative binomial distribution, with the processing batch and sample ID taken as random effects, and the number of UMIs and percentage of reads mapped to mitochondrial genes included as fixed effects. GSEA was performed using the R package fgsea [22]. The results were corrected for multiple comparisons using the Benjamini-Hochberg method [23].

#### Generation of spatial transcriptomics (ST) data

Human kidney biopsies (lupus nephritis and healthy controls) from Brigham and Women’s Hospital were prepared for Xenium spatial transcriptomics. A custom probe set designed to distinguish between all myeloid clusters identified in the scRNA-seq data was used with the manufacturer’s multi-tissue panel. Probe hybridization captured spatially resolved gene expression. Sections were imaged using the Xenium platform, then H&E stained for alignment of gene expression patterns with tissue morphology. Renal compartments were annotated by a board-certified renal pathologist (RS).

#### ST data analysis

Spatially resolved gene expression data were analyzed with Seurat (v5.0) [24] and Xenium Explorer (v3.1). Initial filtering removed cells in the lowest 10% of UMIs and gene counts. Data underwent normalization, feature selection, scaling, dimensional reduction, and batch correction using Harmony [15]. Low-resolution clustering of ST data using Seurat was first applied in order to identify immune and non-immune cells. A similar approach was then used to identify the main immune cell lineages – myeloid cells, T/NK cells and B/plasma cells. Cells identified as myeloid cells were then mapped to the scRNA-seq clusters of myeloid cells, using the reference mapping functionality in Seurat; for the identification of PECs and other tissue non-immune cells, the cells were mapped to the KPMP reference dataset [25]. To validate the mapping, we compared the expression of marker genes separating the scRNA-seq clusters, as found using the scRNA-seq data, in the scRNA-seq and ST datasets. While the final analysis included all myeloid cells in the ST data regardless of the mapping confidence score, we did run a sensitivity analysis, checking the effect of keeping only cells with a mapping confidence score higher than some threshold (several different thresholds were tested); this did not change the reported results in any major way, suggesting they are robust.

#### Identifying preferential positioning in specific kidney region types in ST data

To determine if the cells mapped to each myeloid scRNA-seq cluster demonstrate a statistically-significant preference to occupy a specific type of region of the kidney (glomeruli, inflammatory aggregates in the TI etc), we took the following approach: for each cluster, we first calculated the fraction of cells in each region type. We then compared these fractions to those expected under a null hypothesis stating that no positional preference exists for the cluster; under this null hypothesis, the expected fraction of cells per region type is calculated empirically from the myeloid cells in the ST data without taking into account the identity of their mapped cluster (i.e., assuming that all clusters have the same spatial distribution across region types). P-values were calculated using a binomial distribution, then corrected for multiple comparisons using the Benjamini-Hochberg method [23], taking into account all myeloid clusters and all compartments. Furthermore, given the relatively small number of patients with ST data, we took extra care to avoid reporting results driven by a single patient; to this end, we included only results that were found to have FDR ≤ 0.05 in at least two separate patients, when analyzed individually.

#### Identifying statistically-significant co-localization of monocyte/macrophage clusters in the ST data

To identify pairs of clusters that demonstrate a higher-than-expected co-localization in the kidney ST data, we performed the following steps: for each cluster of monocytes/macrophages (clusters falling under the categories of monocytes, differentiated monocytes, RMs, DMacs in Fig. 1), we counted the number of neighboring cells from each cluster of monocytes/macrophages. We then compared this number to that expected under a null hypothesis according to which there is no preference for co-localization; under this null hypothesis, the expected fraction of neighbors from each monocyte/macrophage cluster is equal to its general fraction out of all monocytes/macrophages (i.e. without knowledge of its position in the kidney). P-values were calculated using a binomial distribution, then corrected for multiple comparisons using the Benjamini-Hochberg method [23], taking into account all pairs of monocyte/macrophage clusters. As done above, we included only results that were found to have FDR ≤ 0.05 in at least two patients, when analyzed individually. Note that this calculation is based on a distance threshold defining cellular adjacency; in the text, we reported results using a threshold of 30μm, and verified their robustness by checking distance thresholds between 20μm and 50μm.

#### Inference of differentiation trajectories

Trajectory analysis using the scRNA-seq data was done using PAGA [26]; we chose this tool because it performs well in benchmark tests [27] and does not restrict the structure of the resulting transitions graph, making it suitable for an analysis allowing for multiple source states, as was the case here. To allow edges to exit all possible origin states (classical monocytes, nonclassical monocytes, Lyve1^+^ RMs and Lyve1^−^RMs) we kept all PAGA edges with a confidence score higher or equal to 0.825 – the highest score existing the Lyve1^−^ RMs (for the other 3 populations, edges with a higher confidence score were available); this was justified by the fact that for certain DMac clusters (e.g. M7), the percentage of cells most similar to Lyve1^−^ RMs, as calculated by reference mapping, was relatively large, i.e. transitions out of this RM population are plausible. Next, we identified the shortest paths from each of the 4 origin states to each DMac cluster. We then checked which of the edges on these trajectories connect clusters that demonstrated statistically significant co-localization in the ST data.

#### Cell cultures

CD14 monocytes were isolated using negative selection (Miltenyi) from three healthy donors, pooled, and >90% purity was verified by flow cytometry. A total of 50,000 CD14 monocytes per well were aliquoted in a 96-well format. The cells were initially treated with 50 ng/mL M-CSF (Peprotech Cat. # 300-25) in standard media (RPMI, 10% FBS, Penicillin, Streptomycin) for five days. At this point, the media was replaced, maintaining M-CSF while introducing 50 ng/mL of TGF-β (Peprotech Cat. # 100-21), no stimulus, or IFN-γ (Peprotech Cat. # 300-02) for an additional two days. Following treatment, the cells were hashed according to the manufacturer’s protocol and prepared for 3’ single-cell RNA sequencing using the 10X Chromium platform.

#### Ligand-receptor analysis for PECs

Differential expression analysis for PECs, comparing patients with gCI=0 to LD controls was performed as described above. We next looked for receptors found to be differentially expressed (FDR ≤ 0.05), whose ligands were previously shown to induce the activation, proliferation or migration of epithelial or endothelial cells. Ligand-receptor pairs were based on CellPhoneDB v4.1.0 [28]. As this dataset contains information about the subunits of each receptor, a receptor was considered differentially expressed if at least one of the genes coding these subunits was differentially expressed. Furthermore, we required all subunits of the receptor to be expressed by at least 30% of the PECs for it to be considered. We then focused on receptors whose ligands are expressed in a relatively high level in at least one of the clusters of differentiated monocytes and Dmacs that demonstrate statistically significant co-localization with PECs.

#### Identifying myeloid clusters enriched in the vicinity of PECs in the ST data

We computed the actual probability of cells from each myeloid cluster to be within some distance *r* of PECs, defining a proximity threshold, and compared it to the probability expected if immune cells were spread in the tissue uniformly (the results reported in the manuscript were found to be robust to the choice of *r* within the range 10-100μm). To do this, we first computed the average number of cells within *r* μm of PECs, *m*, based on the actual empirical data. Under the null hypothesis, i.e. if immune cells were spread uniformly in the tissue, then for each myeloid cluster *j*, the probability that a cell drawn by chance from the proximity of a PEC would be of that cluster is equal to the proportion of that cluster out of all cells in the tissue, *f*_*j*_. Thus, under the null hypothesis, the probability that at least one of the cells in the proximity of a PEC would be of cluster *j* is equal to *P*_0*j*_ = 1 − [1 − *f*_*j*_] . To compute the p-value, we compared this probability to the one actually observed using a Binomial test. The p-values across all myeloid clusters were then corrected for multiple comparisons using the Benjamini-Hochberg method [23].

#### Defining PEC densities

For each PEC, we counted the number of PECs within 30μm of it. We defined low PEC density to be less than 5 adjacent PECs; intermediate density – 5 or more, less than 8 adjacent PECs; and high density – 8 or more adjacent PECs. These thresholds were determined arbitrarily based on visual inspection of the ST images; we found that the reported results are robust to a wide range of their values.

---

**Supplementary Figure 1.**
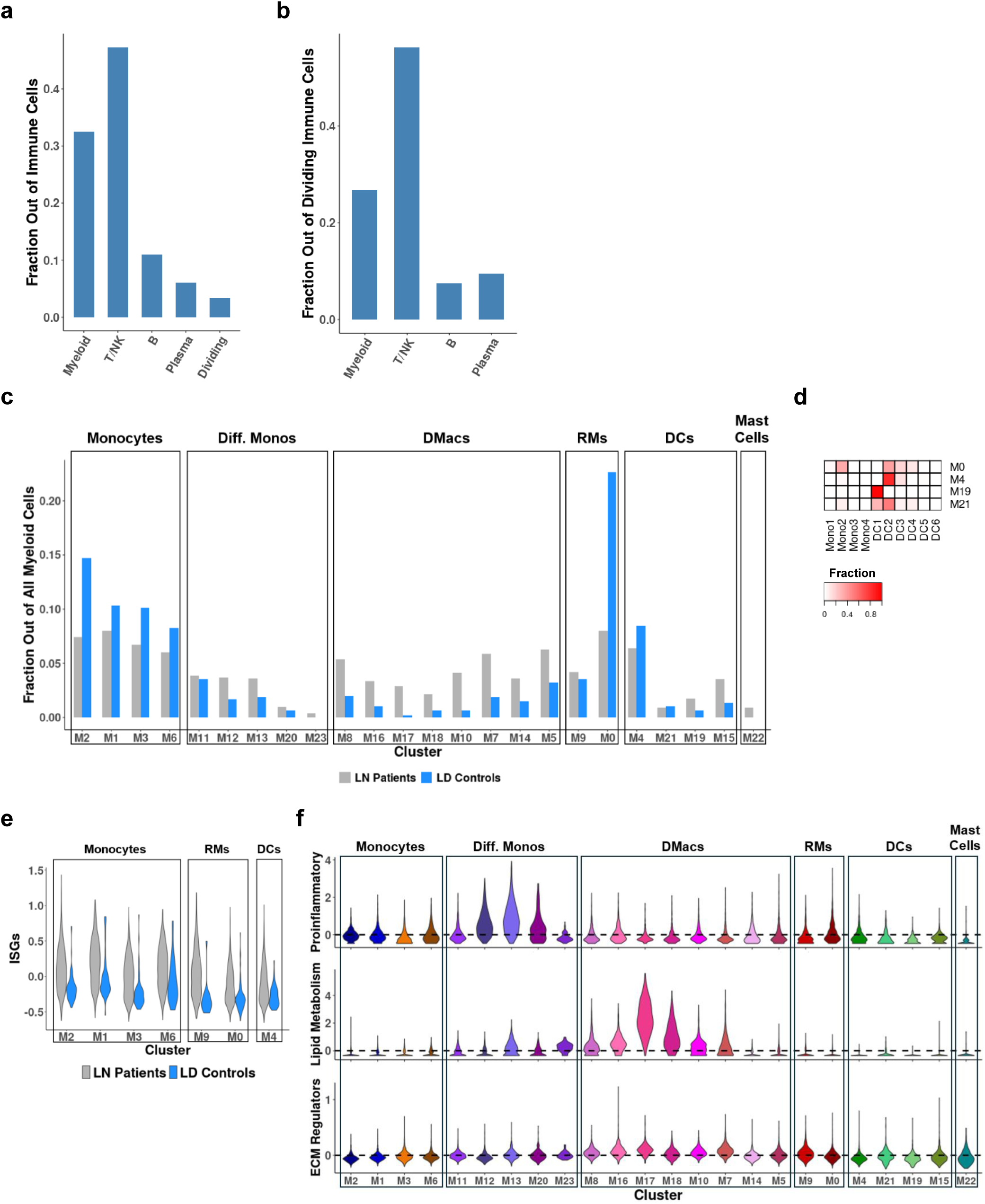
(a) The fraction of cells per lineage, out of all immune cells. (b) The fraction of dividing cells per lineage, out of all dividing cells. (c) The fraction of cells included in each myeloid cell cluster, in LN patients and in LD controls. (d) The fraction of cells mapped to each reference class in Villani et al, 2017, specified for myeloid cell clusters predominately mapped to the “DC” class in Stewart et al., 2019. (e) The distribution of the “interferon score”, computed per cell as the average of scaled expression of ISGs, across myeloid cell clusters, comparing LN patients and LD controls. Shown are only clusters abundant in LD controls. (f) The distribution of three composite scores, computed per cell as the average of scaled expression of a gene set of interest, across myeloid cell clusters. Only cells from LN patients were taken into account. The dashed line represents the average across all cells from LN patients, regardless of cluster. Top: a composite score computed based on proinflammatory cytokines and chemokines. Middle: a composite score computed based on genes associated with lipid metabolism. Bottom, a composite score computed using genes previously identified as ECM regulators.

**Supplementary Figure 2.**
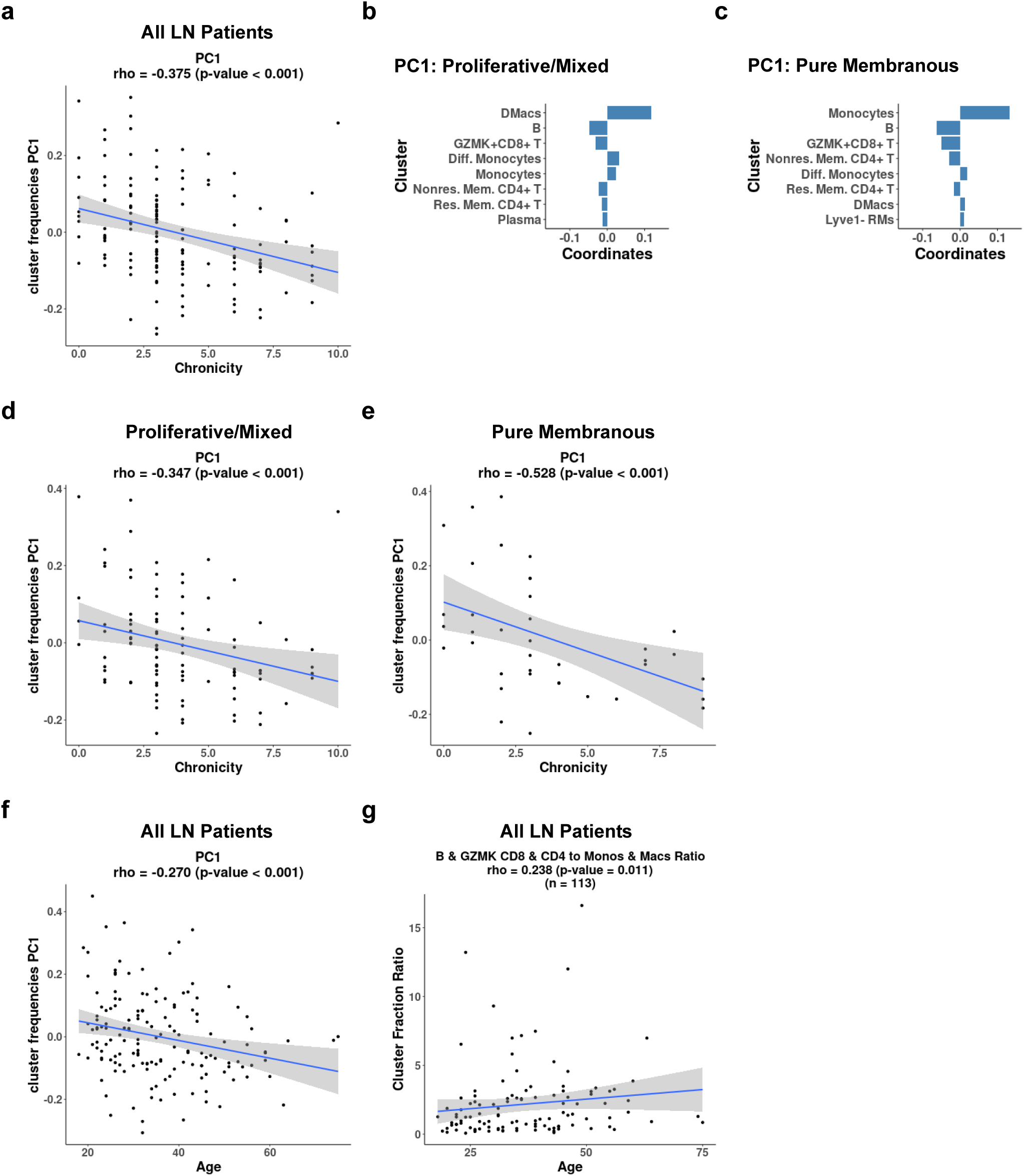
The results of using PCA to analyze variability in cluster frequencies. In all panels, only patients with at least 20 immune cells were considered; the reported results are robust to the choice of this threshold within the 10-100 cells range. Panels (a), (d), (e), (f) and (g) report the Spearman correlation and its associated p-value. (a) The correlation between the NIH chronicity index and PC1, taking into account all LN patients. (b) The top contributing clusters to PC1, in PCA considering only proliferative/mixed LN patients. (c) The top contributing clusters to PC1, in PCA considering only pure membranous LN patients. (d) The correlation between the NIH chronicity index and PC1, in proliferative/mixed LN patients. (e) The correlation between the NIH chronicity index and PC1, in pure membranous LN patients. (f) The correlation between the age of the patients, at the time the biopsy was obtained, and PC1, considering all LN patients. (g) The correlation between the age of the patients, at the time of biopsy acquisition, and the ratio between the frequencies of selected lymphoid and myeloid clusters, as described in the text, considering all LN patients.

**Supplementary Figure 3.**
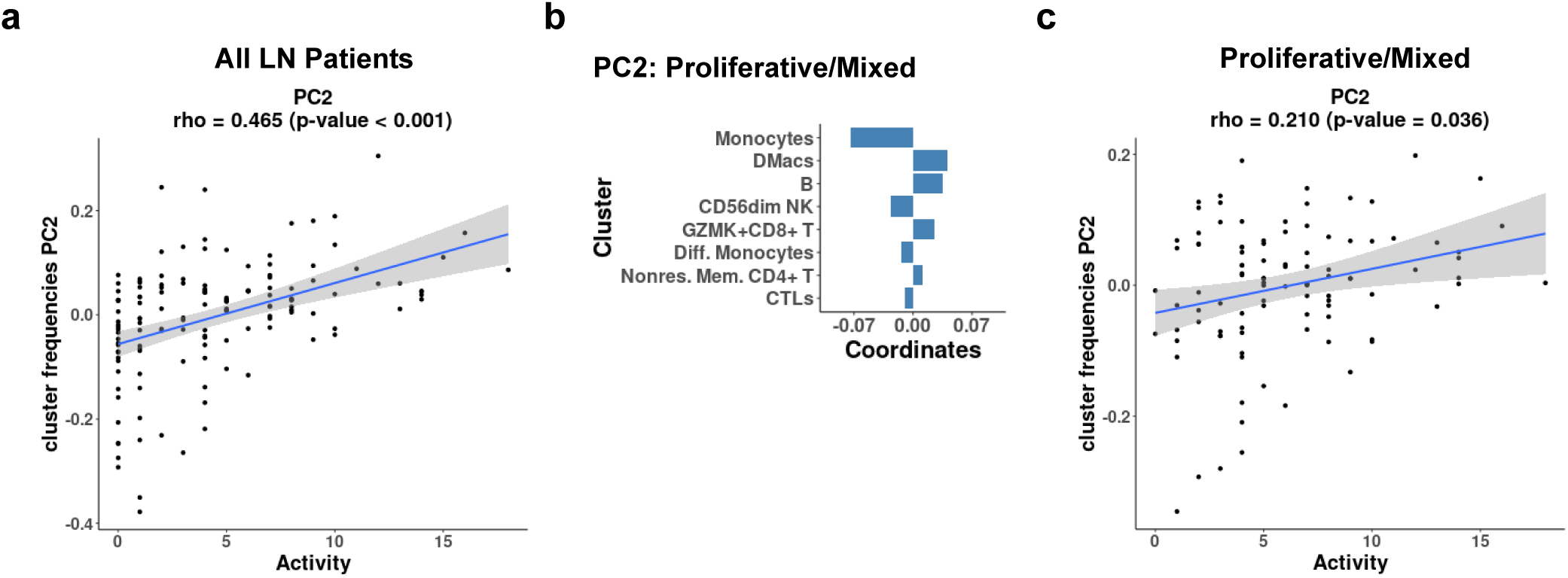
The results of using PCA to analyze variability in cluster frequencies. In all panels, only patients with at least 20 immune cells were considered; the reported results are robust to the choice of this threshold within the 10-100 cells range. Panels (a) and (c) report the Spearman correlation and its associated p-value. (a) The correlation between the NIH activity index and PC2, taking into account all LN patients. (b) The top contributing clusters to PC2, in PCA considering only proliferative/mixed LN patients. (c) The correlation between the NIH activity index and PC2, taking into account only proliferative/mixed LN patients.

**Supplementary Figure 4.**
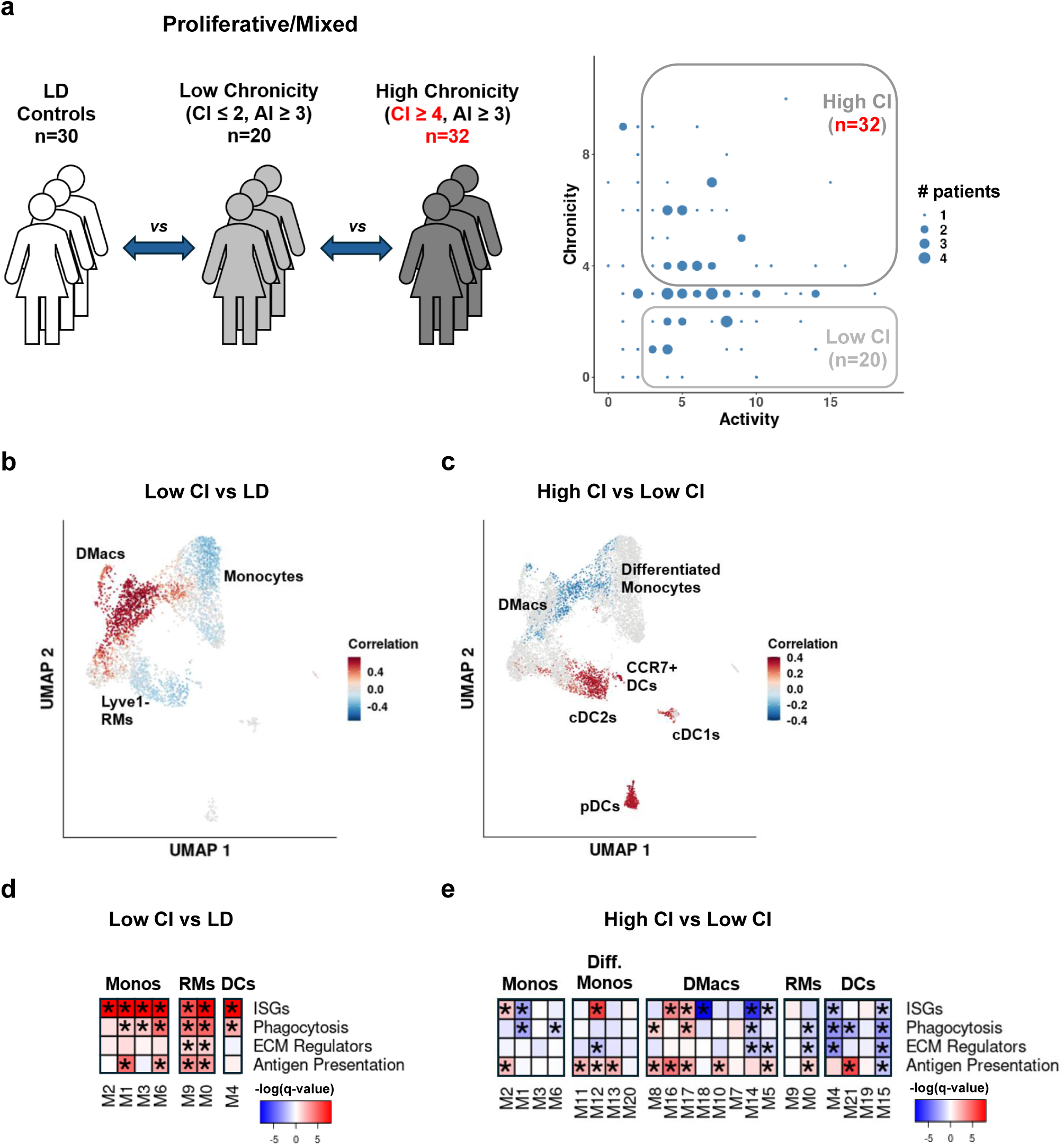
An identical analysis to that reported in Figure 3, while setting the threshold defining the high CI group to 4 instead of 5. Differences from Figure 3 are marked in red.

**Supplementary Figure 5.**
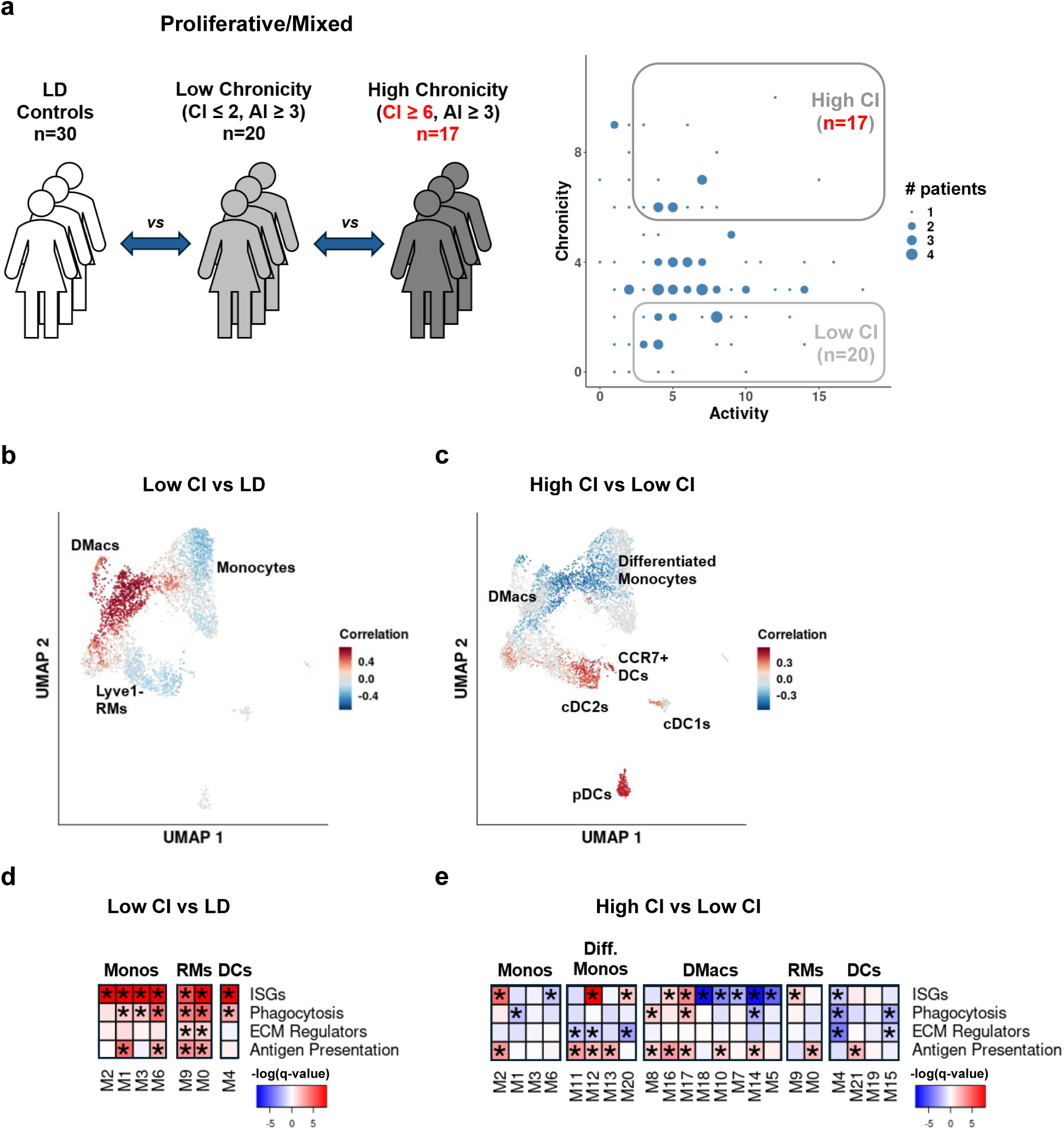
An identical analysis to that reported in Figure 3, while setting the threshold defining the high CI group to 6 instead of 5. Differences from Figure 3 are marked in red.

**Supplementary Figure 6.**
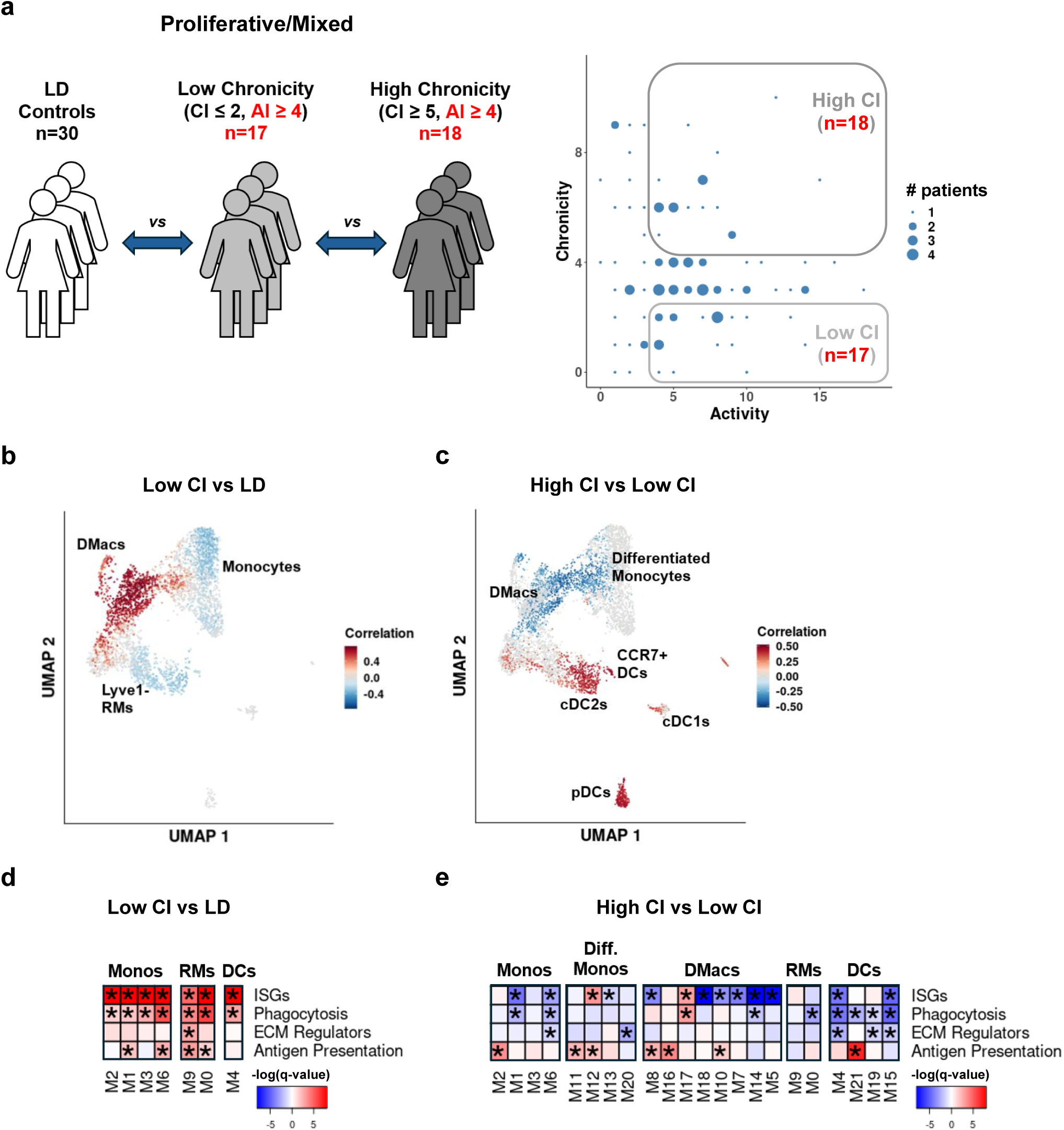
An identical analysis to that reported in Figure 3, while setting the threshold defining basal AI to 4 instead of 3. Differences from Figure 3 are marked in red.

**Supplementary Figure 7.**
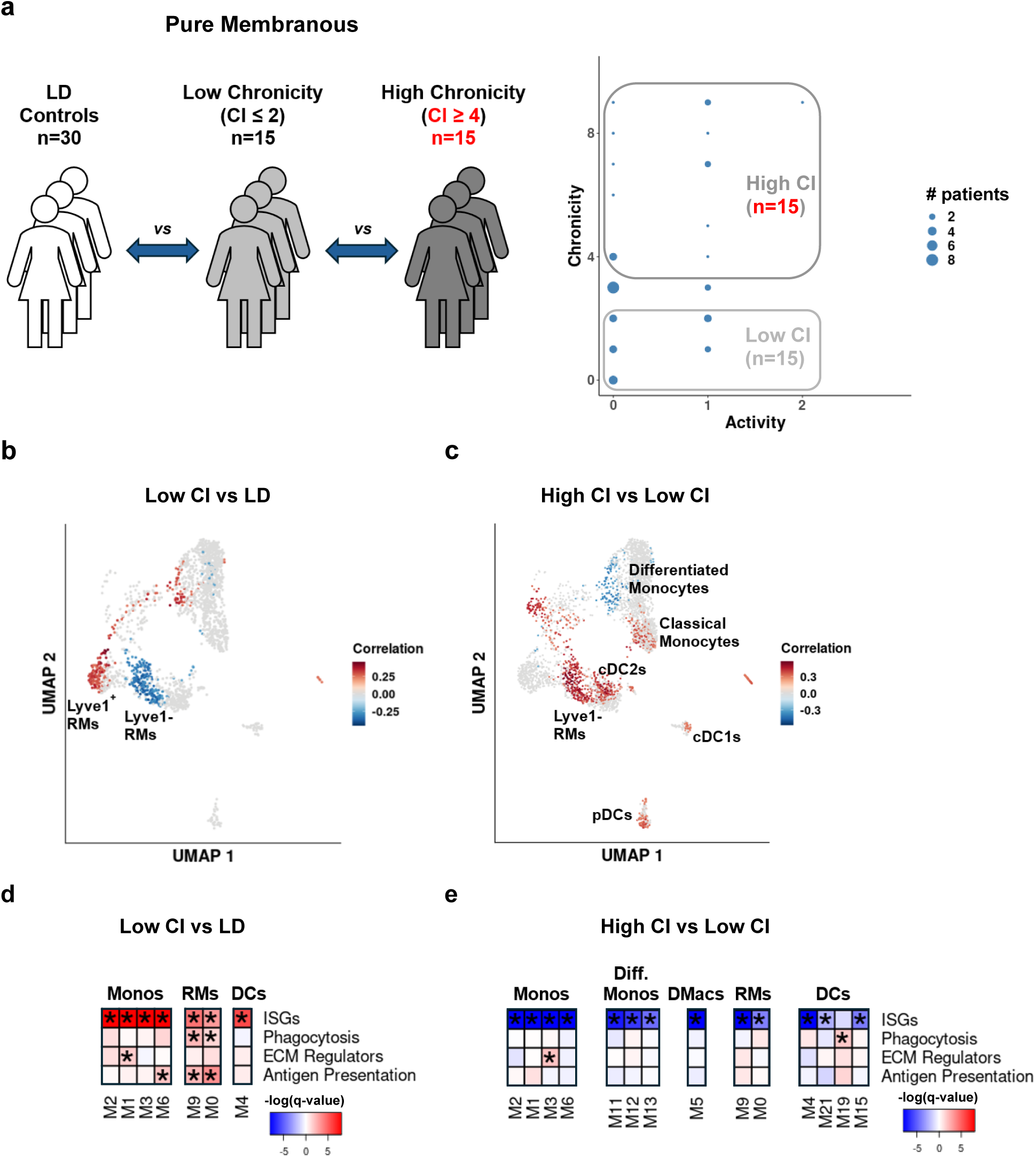
An identical analysis to that reported in Figure 4, while setting the threshold defining the high CI group to 4 instead of 5. Differences from Figure 4 are marked in red.

**Supplementary Figure 8.**
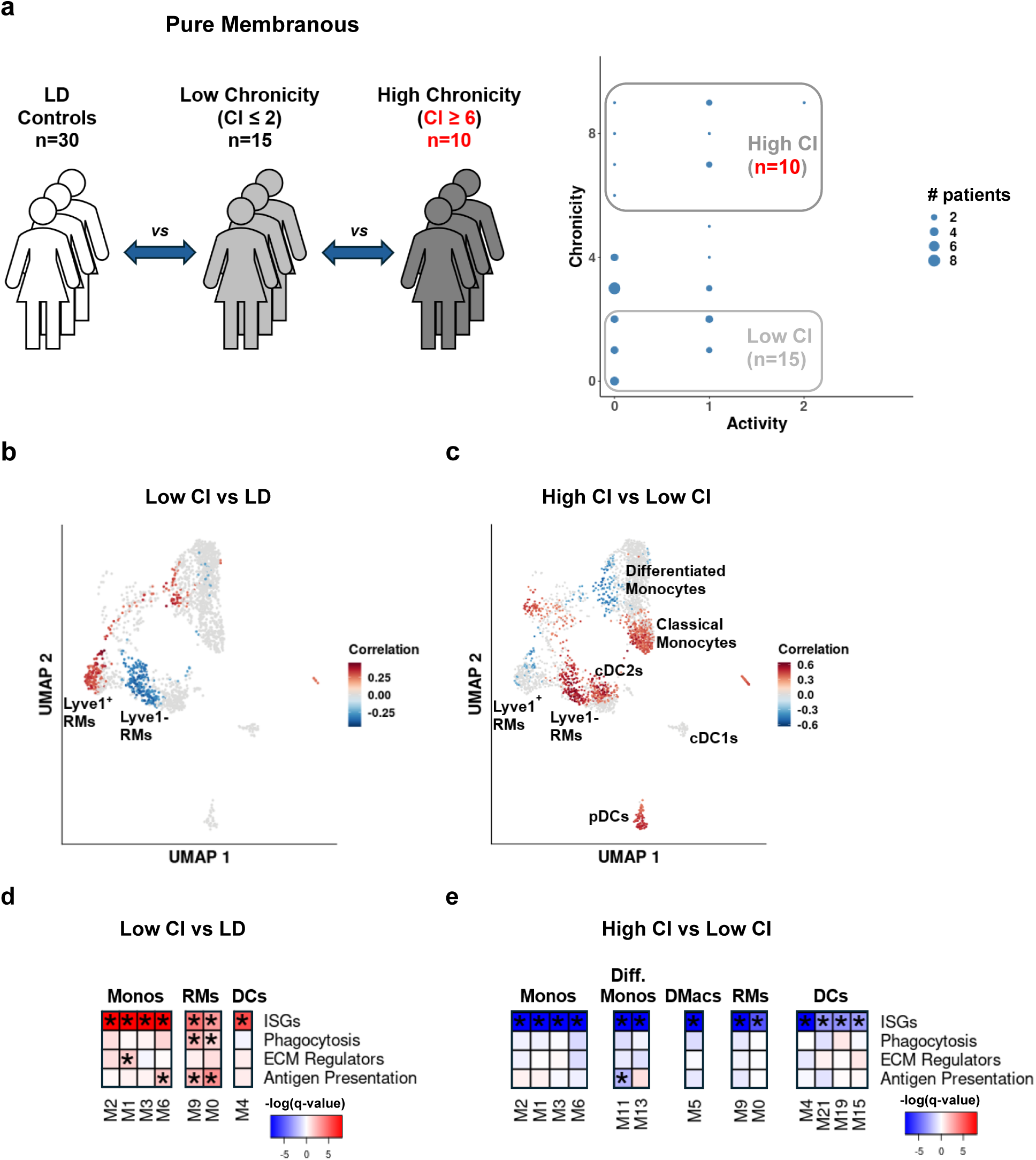
An identical analysis to that reported in Figure 4, while setting the threshold defining the high CI group to 6 instead of 5. Differences from Figure 4 are marked in red.

**Supplementary Figure 9.**
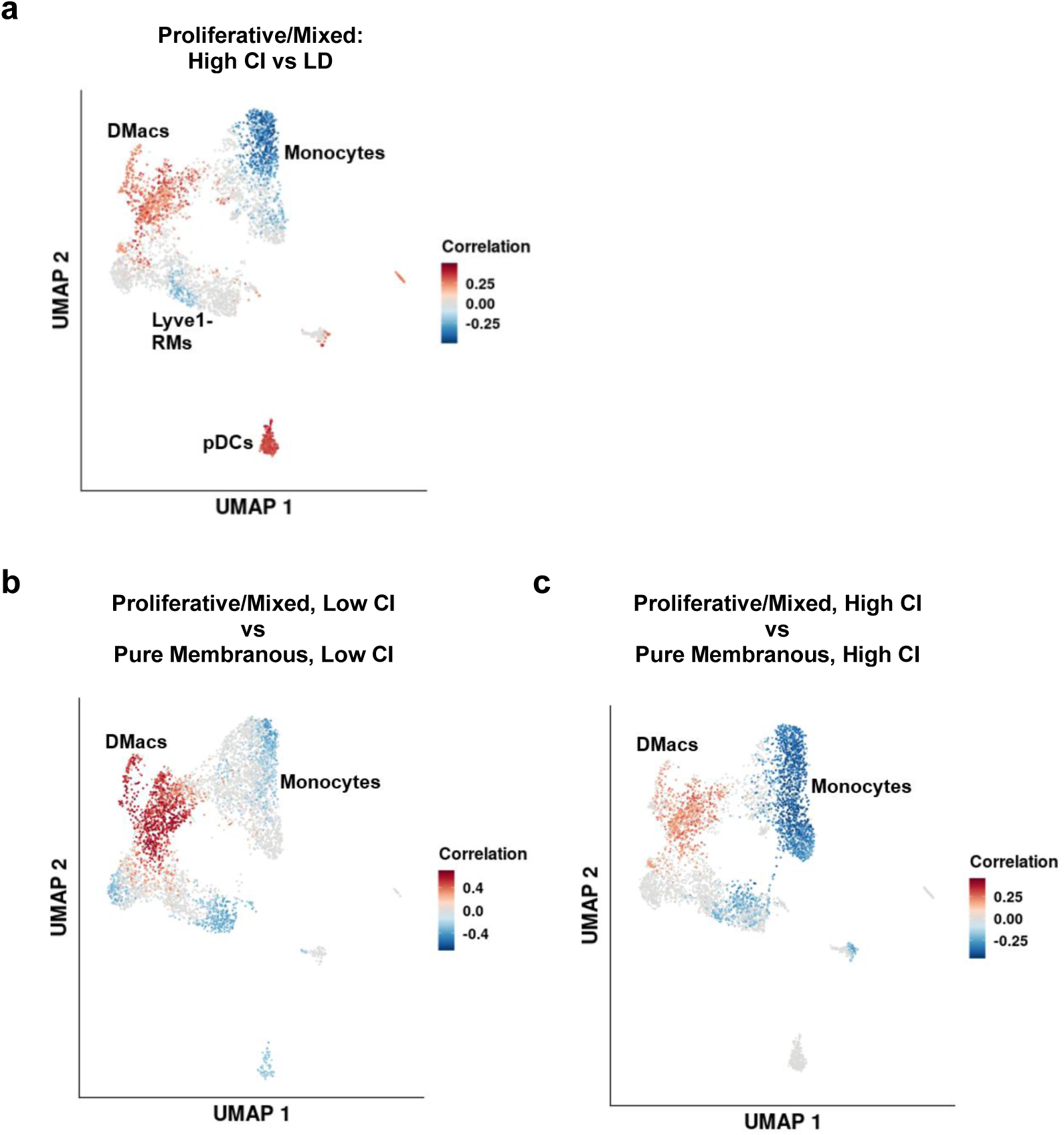
(a) The results of running CNA, comparing the proliferative/mixed LN patients with High CI to LD controls. Cell populations expanded in LN patients are colored in red, those depleted in blue. (b) The results of CNA, comparing the Low CI proliferative/mixed patients to the Low CI pure membranous patients. Populations expanded in the proliferative/mixed patients are colored in red; those depleted are colored in blue. (c) The results of CNA, comparing the High CI proliferative/mixed patients to the High CI pure membranous patients. Populations expanded in the proliferative/mixed patients are colored in red; those depleted are colored in blue. In all panels, only correlations found to be statistically-significant after correction for multiple comparisons (FDR ≤ 0.1) are colored.

**Supplementary Figure 10.**
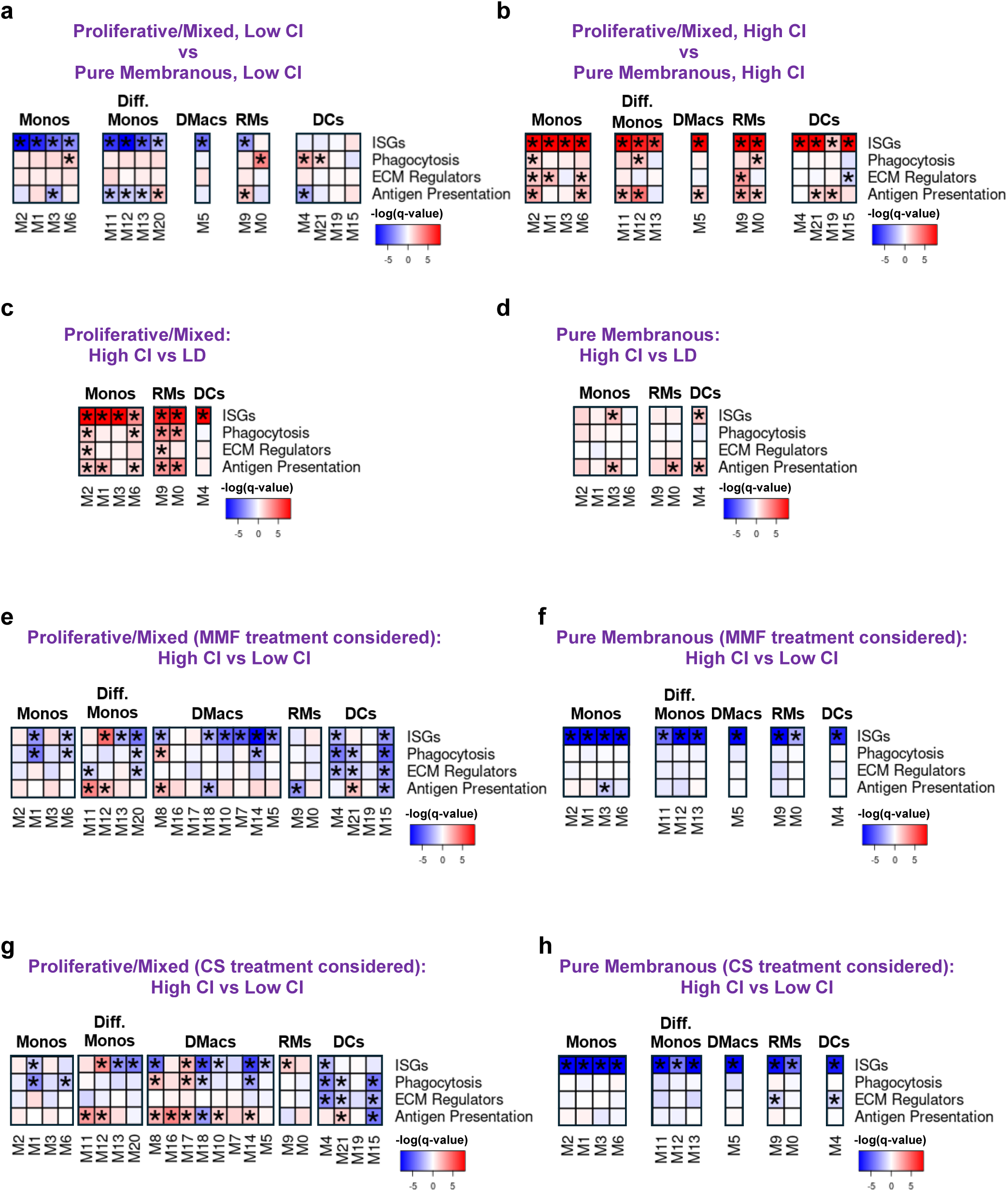
All panels report the results of running GSEA following DE analysis per cluster. Asterisks mark FDR ≤ 0.05. The color of each box was determined by the significance of the enrichment (darker shades mean more significant enrichment), and the sign of the enrichment, as specified below. (a) Comparison of Low CI proliferative/mixed patients and Low CI pure membranous patients. Red – higher in proliferative/mixed LN, blue denotes the opposite. (b) Comparing the High CI proliferative/mixed patients to the High CI pure membranous patients. Red represents upregulation in proliferative/mixed LN, blue – downregulation. (c) Comparison of High CI proliferative/mixed patients and LD controls. Red designates higher values in LN patients, blue – higher values in LD controls. (d) Comparison of High CI pure membranous patients and LD controls. Red – higher values in LN patients, blue – higher values in LD controls. (e) Comparison of High CI and Low CI proliferative/mixed patients, using a model that includes an indicator variable denoting treatment with MMF. Red – upregulated in High CI group; blue – downregulated. (f) Comparison of High CI and Low CI pure membranous patients, using a model that includes an indicator variable denoting treatment with MMF. Red represents upregulation in the High CI group, blue designates downregulation. (g) Same as (e), with a flag denoting treatment with high-dose (≥ 30mg) corticosteroids. (h) Same as (f), with a flag denoting treatment with high-dose (≥ 30mg) corticosteroids.

**Supplementary Figure 11.**
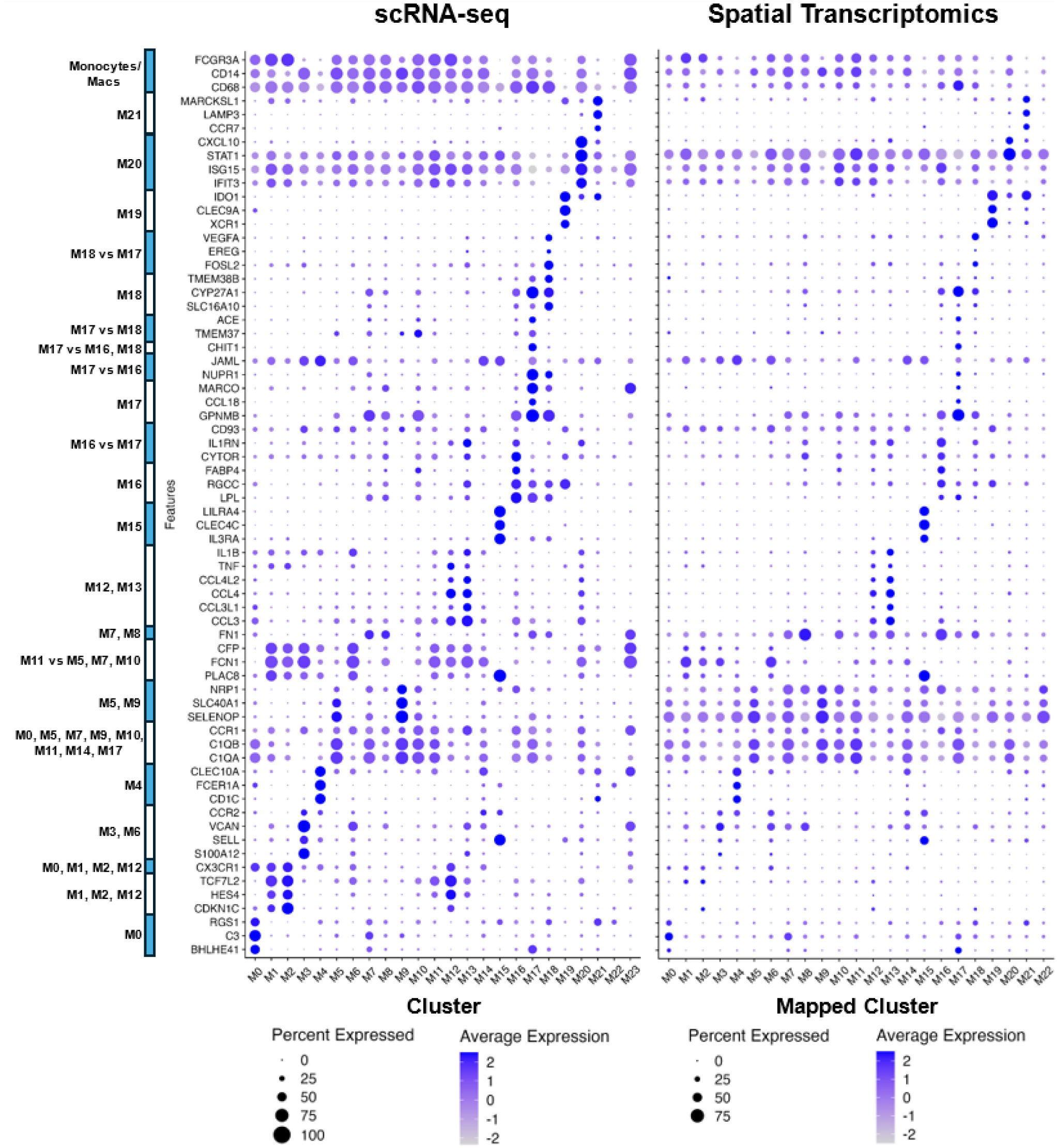
Comparison of the expression of selected genes in the scRNA-seq dataset (left) and the spatial transcriptomics dataset (right). Cells in the scRNA-seq dataset were grouped based on their cluster; cells in the spatial transcriptomics dataset were grouped based on the scRNA-seq cluster they were mapped to. The genes included were chosen based on their potential utility in distinguishing between the clusters identified in the scRNA-seq dataset, as specified on the left. We note, however, that all genes in the spatial transcriptomics data were used for the mapping of cells to the scRNA-seq clusters.

**Supplementary Figure 12.**
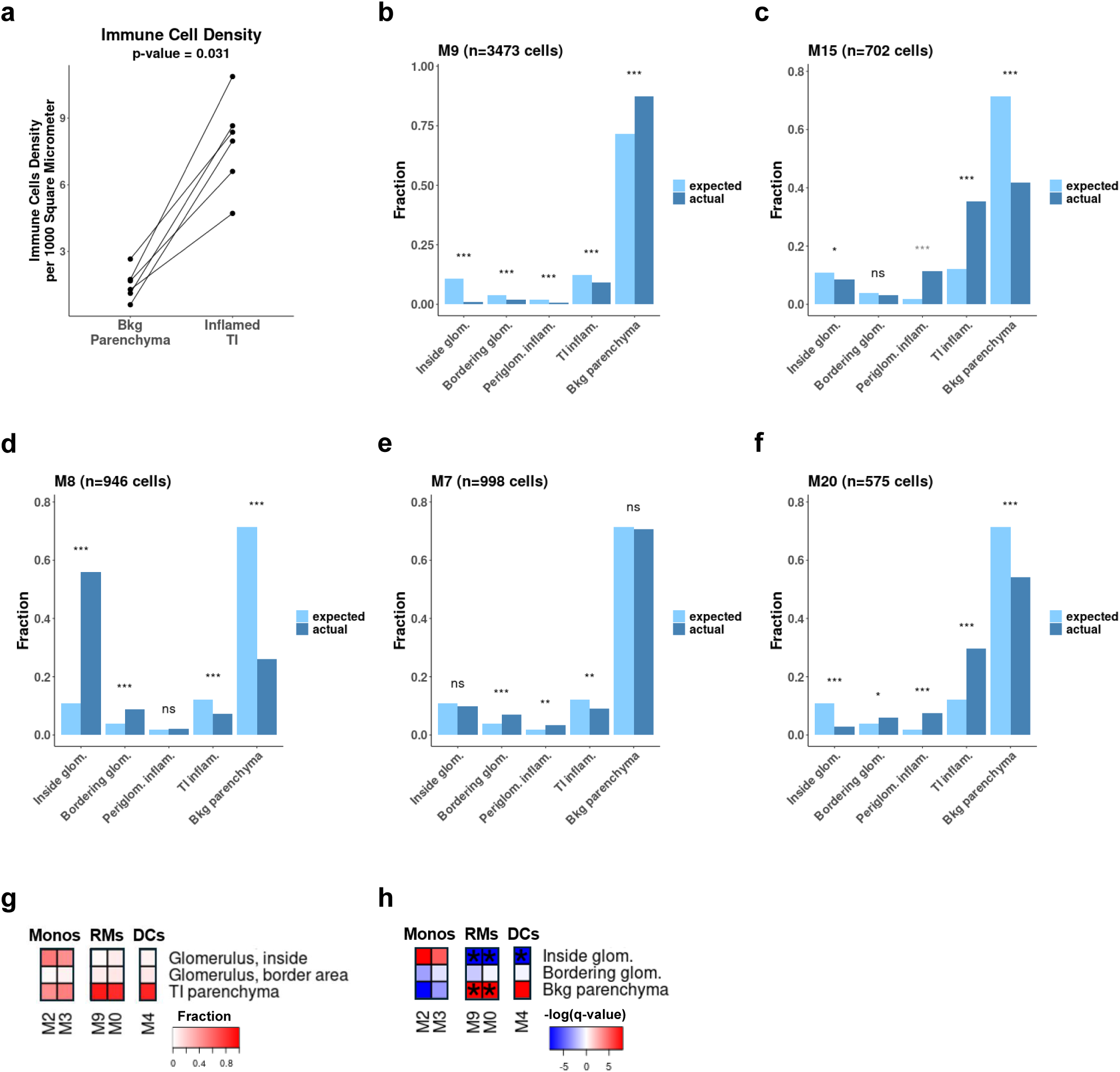
(a) The density of immune cells, per 1000 square micrometer, comparing regions of inflamed TI vs the normal background parenchyma. Each point represents a LN patient; lines connect measurements corresponding to the same patient. The p-value was calculated using the paired Wilcoxon signed rank test. (b-f) The spatial distribution in LN kidneys of the cells of individual clusters. Five clusters are shown as an example. For each cluster, dark blue designates the actual fraction of cells in each type of region of the tissue (glomeruli, inflamed TI, etc.); light blue represents the expected fraction, if the cluster cells did not have a preference to occupy specific region types. *** - FDR ≤ 0.001; ** - FDR ≤ 0.01; * - FDR ≤ 0.05; ns – not significant. Greyed-out asterisks are not supported by the individual analysis of at least two study subjects (here, this occurs only for cluster M15 cells in periglomerular inflammation). (b) The spatial distribution of cluster M9 (Lyve1^−^ RMs). (c) The spatial distribution of cluster M15 (pDCs). (d) The spatial distribution of cluster M8 (proinflammatory-like DMacs). (e) The spatial distribution of cluster M7 (phagocytic-like DMacs). (f) The spatial distribution of cluster M20 (ISG-high monocytes/macrophages). (g) The spatial distribution of myeloid cells in healthy kidneys. Each row represents a specific region type (glomeruli, TI etc.); each column represents a cluster, and shows the fraction of that cluster in each region type (the sum of values per cluster is equal to 1). (h) The statistical significance of positional preferences, in healthy kidneys. Each row represents a specific region type; each column represents a cluster. Heatmap elements are colored based on the magnitude of the adjusted p-values, and the direction of observed deviance from the null distribution assuming no preference: red represents a relative preference of a cluster to occupy a specific region type; blue represents a relative avoidance. Asterisks represent cases with FDR ≤ 0.05, further supported by the individual analysis of both healthy subjects.

**Supplementary Figure 13.**
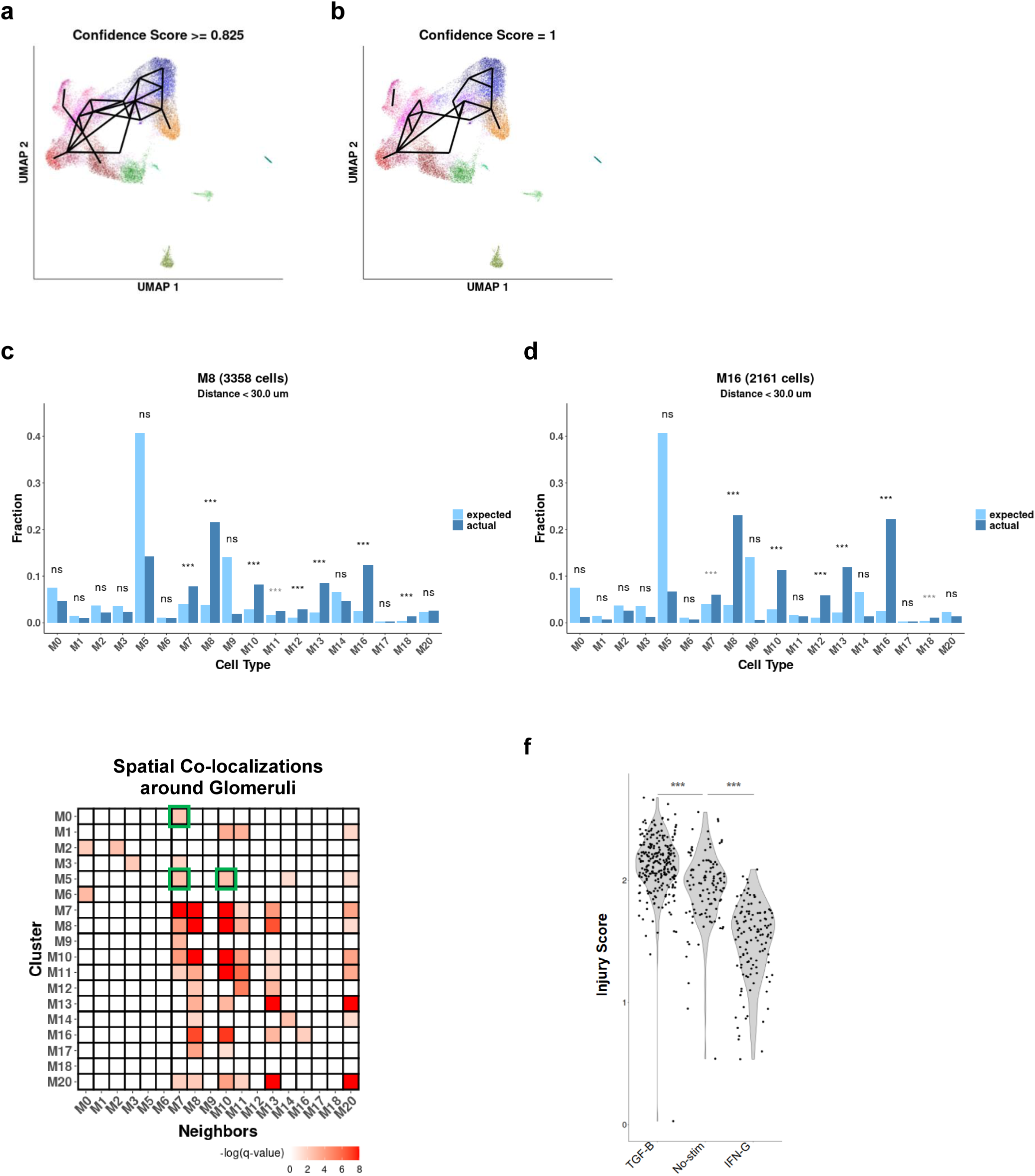
(a) The results of trajectory analysis using PAGA. Shown are edges with a confidence score ≥ 0.825, which is the threshold required to allow transitions out of the Lyve1- RMs (cluster M0). (b) Same as (a), showing edges with highest confidence score possible, 1. (c) The distribution of monocyte & macrophages neighbors of cells in cluster M8. Dark blue designates the observed fraction out of all monocyte/macrophages neighbors; light blue represents the expected fraction, if no preferred co-localization existed. *** - FDR ≤ 0.001; ** - FDR ≤ 0.01; * - FDR ≤ 0.05; ns – not significant. Greyed-out asterisks are not supported by the individual analysis of at least two study subjects. (d) The same as (c), for cells in cluster M16. (e) Statistically significant co-localizations of cluster pairs in LN kidneys, with a focus on monocytes and macrophages in regions of periglomerular inflammation or those bordering glomeruli. Each row pertains to a specific cluster, and shows co-localizations with other clusters of monocytes/macrophages. Colored heatmap elements represent cases with FDR ≤ 0.05. The observed co-localizations of cluster pairs M0-M7, M5-M7 and M5-M10 are highlighted in green. Here, cells are considered adjacent if they are within 30μm of each other; similar results are found for choices of this distance threshold between 30μm and 50μm. (f) The expression of the injury-associated gene program in classical monocytes treated with M-CSF in addition to TGFβ, no stimulus, or IFNγ, as measured by scRNA-seq. P-values were computed using the Mann-Whitney U test. *** - p-value ≤ 0.001.

**Supplementary Figure 14.**
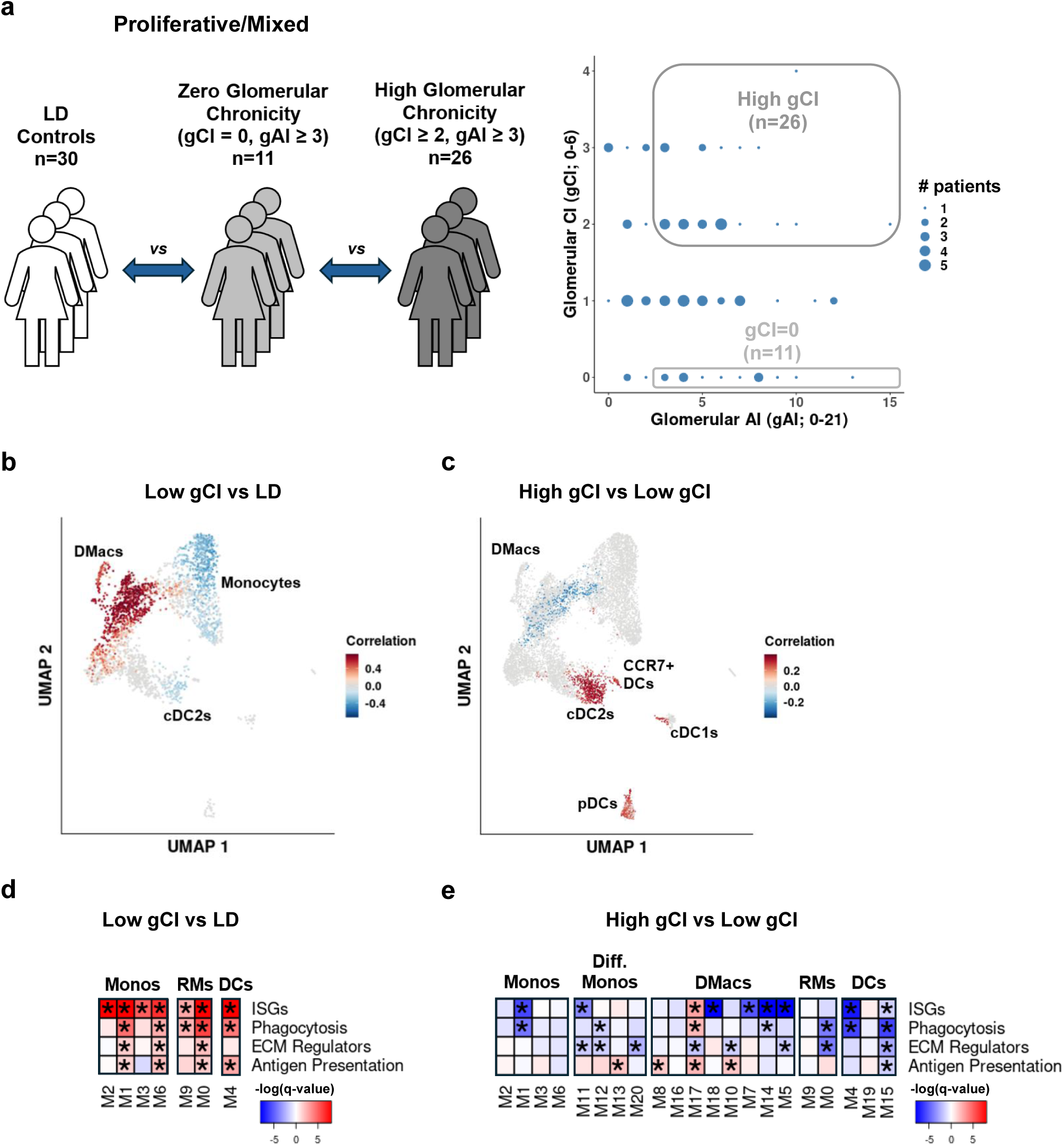
An identical analysis to that reported in Figure 3, while defining patient groups based on the glomerular components of the NIH activity and chronicity indices. For the chronicity, the glomerular components are: global or segmental glomerulosclerosis, fibrous crescents. For the activity, the glomerular components are: endocapillary hypercellularity, neutrophils karyorrhexis, wire loops or hyaline thrombi, fibrinoid necrosis, cellular or fibrocellular crescents.

**Supplementary Figure 15.**
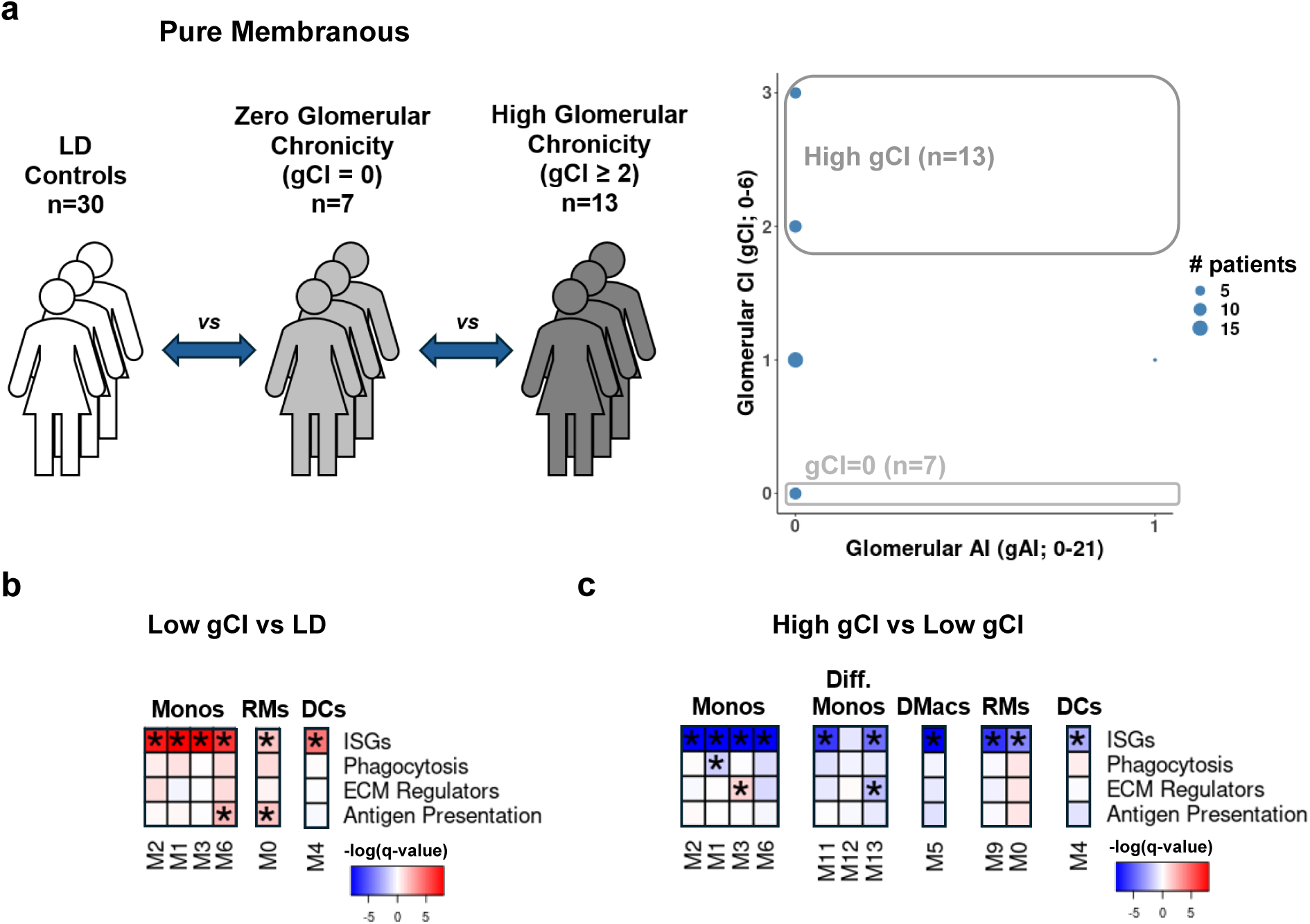
As in Supplementary Figure 14, limiting the analysis to patients with pure membranous LN. Note that in this case, running CNA did not identify significant changes for either of the two comparisons made, and therefore the corresponding UMAP plots are not shown.

**Supplementary Figure 16.**
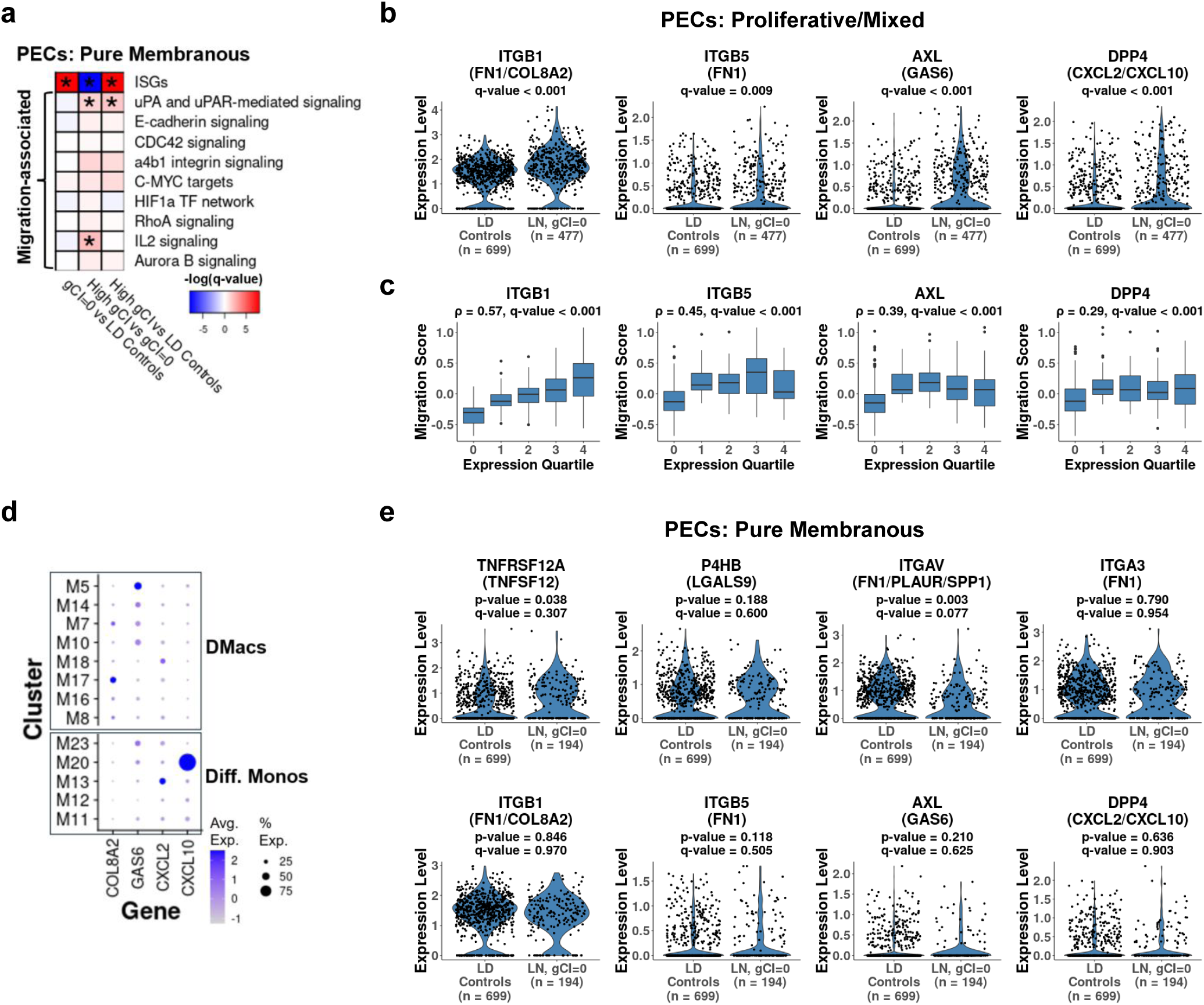
(a) The results of running GSEA following DE analysis for PECs, comparing: (i) the gCI=0 pure membranous patients and LD controls (left; red means upregulation in LN patients); (ii) High gCI and gCI=0 pure membranous patients (middle; red denotes upregulation in the High gCI patients), and (iii) High gCI pure membranous patients and LD controls (right; red represents upregulation in LN patients). Tested were ISGs as well as gene sets previously shown to be associated with epithelial cell migration. Asterisks mark FDR ≤ 0.05. The color of each box was determined by the significance of the enrichment (darker shades mean more significant enrichment) and the sign of the enrichment. (b) Changes in the expression levels in PECs of several receptors, comparing the gCI=0 proliferative/mixed patients and LD controls. For each gene, the FDR-adjusted p-value (q-value) is specified. n denotes the number of cells included in each group. The corresponding ligands are specified in parentheses. (c) The relation between the expression level of each receptor shown in (b) and the migration score, in each PEC. Included are only PECs from gCI=0 proliferative/mixed patients. Cells were grouped into bins based on the expression level of the receptor shown in each panel; one bin includes cells with an expression value of 0, while the rest of the bins correspond to the quartiles of the expression distribution of the receptor, taking into account only the positive expression values. For each panel, the Spearman correlation ρ and its associated p-value are reported, taking into account all PECs from gCI=0 proliferative/mixed LN patients. (d) The expression, in differentiated monocyte and DMac clusters, of ligands binding the receptors appearing in (b). (e) The expression of the receptors appearing in Figure 6b and in panel (b) here, comparing gCI=0 pure membranous patients and LD controls. For each receptor, the p-value and the FDR-adjusted p-value (the q-value), as calculated in DE analysis taking into account all genes, are reported. The corresponding ligands are specified in parentheses.

**Supplementary Figure 17.**
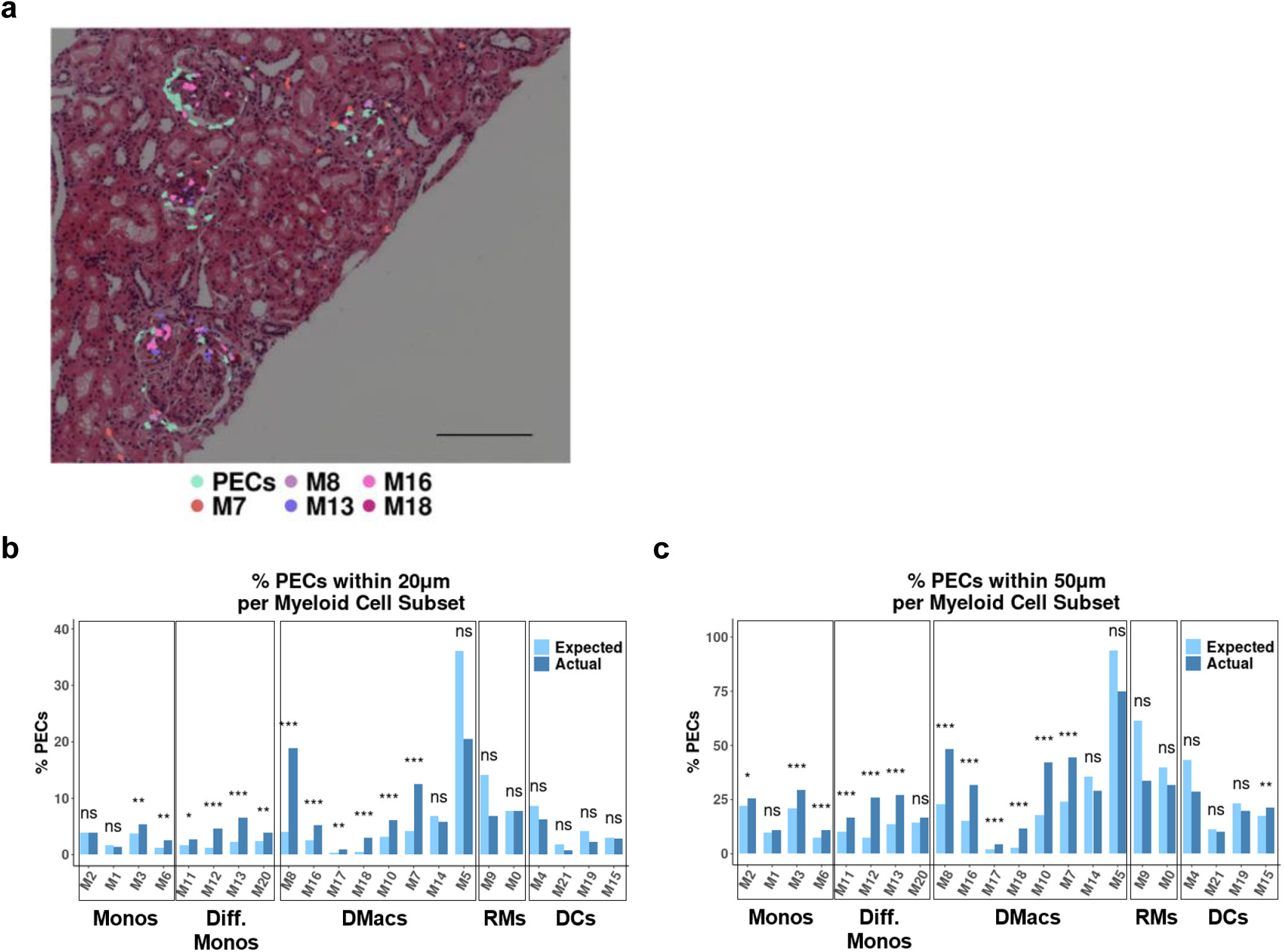
(a) A representative example showing the co-localization of PECs and selected myeloid cell subsets, from the spatial transcriptomics dataset analyzed. Cell positions are shown overlaid on H&E staining. Scale bar denotes 200μm. (b) The percentage of PECs adjacent to each subset of myeloid cells, within a distance of 20μm, in the spatial transcriptomics dataset analyzed. Dark blue shows the actual percentage, light blue the percentage expected if immune cells were distributed uniformly in the tissue. *** - FDR < 0.001; ** - FDR < 0.01; * - FDR < 0.05; ns – not significant. (c) The same as (b), for a distance threshold of 50μm.

**Supplementary Figure 18.**
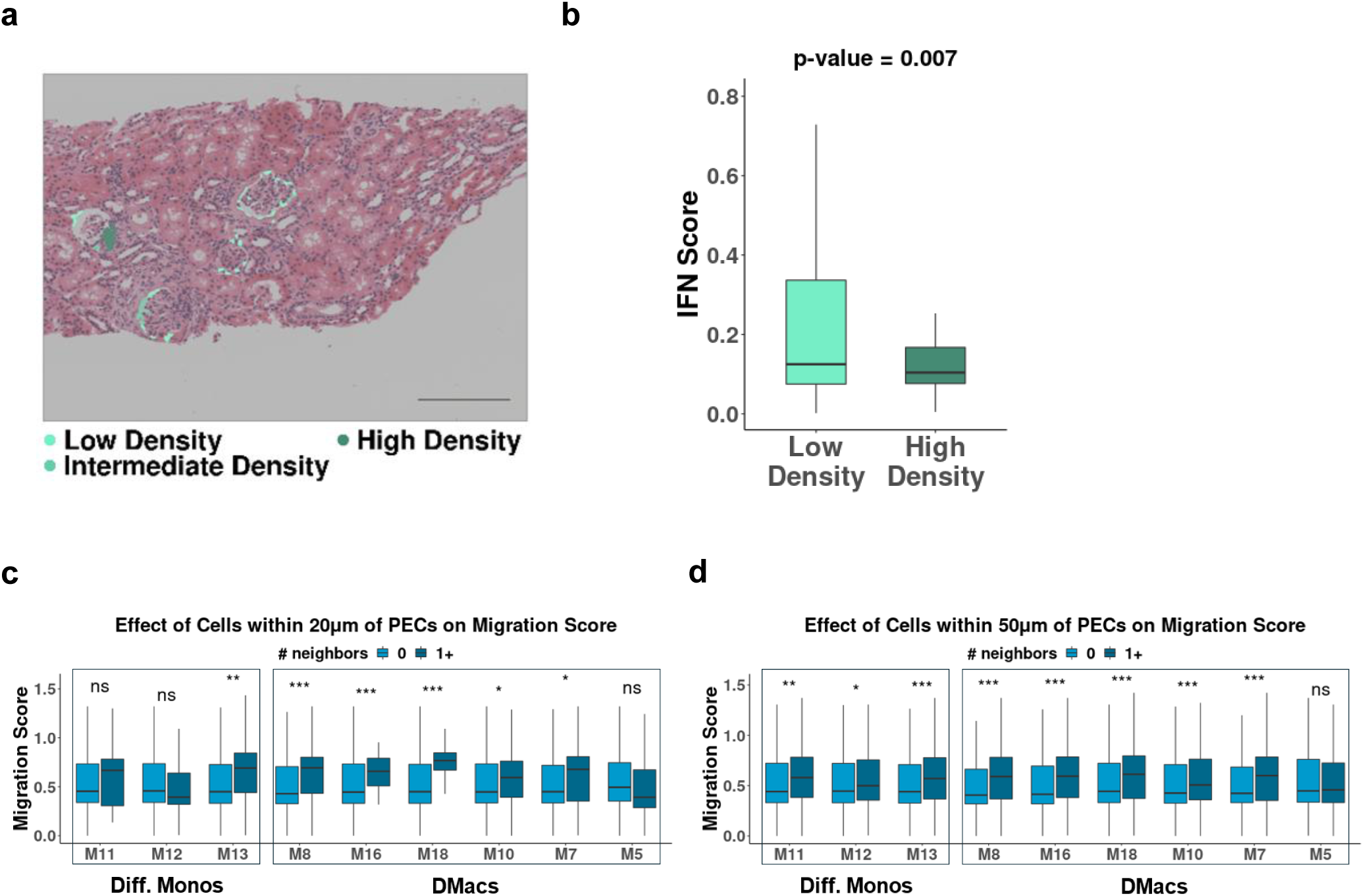
(a) A representative example of PEC aggregates, in a slide taken from the spatial transcriptomics dataset analyzed. Shown are PECs, colored by the number of PECs in their immediate neighborhoods (with a threshold set arbitrarily to 30μm), and overlaid with . Light green – PECs from low density neighborhoods (less than 5 adjacent PECs); green – intermediate density neighborhoods (5 or more, less than 8); dark green – high density neighborhoods (PEC aggregates; 8 or more adjacent PECs). The reported results were found to be robust to the actual choice of these thresholds. Scale bar denotes 200μm. (b) The interferon score, defined by the expression of selected ISGs, in PECs found in low-density and high-density environments, in the spatial transcriptomics dataset analyzed. The p-value was computed using the Mann-Whitney U test. (c) The effect of neighboring cells from selected myeloid clusters on the migration score in PECs. Here, cells are considered adjacent if they are within 20μm of each other. For each myeloid cluster, light blue designates the case of PECs with no neighbors of that cluster, and dark blue represents the case where there is at least one such neighbor. P-values were computed using the Mann-Whitney U test, and corrected for multiple comparisons using the Benjamini-Hochberg method. *** - FDR < 0.001; ** - FDR < 0.01; ns – not significant. (d) The same as (c), for a distance threshold of 50μm.

**Supplementary Figure 19.**
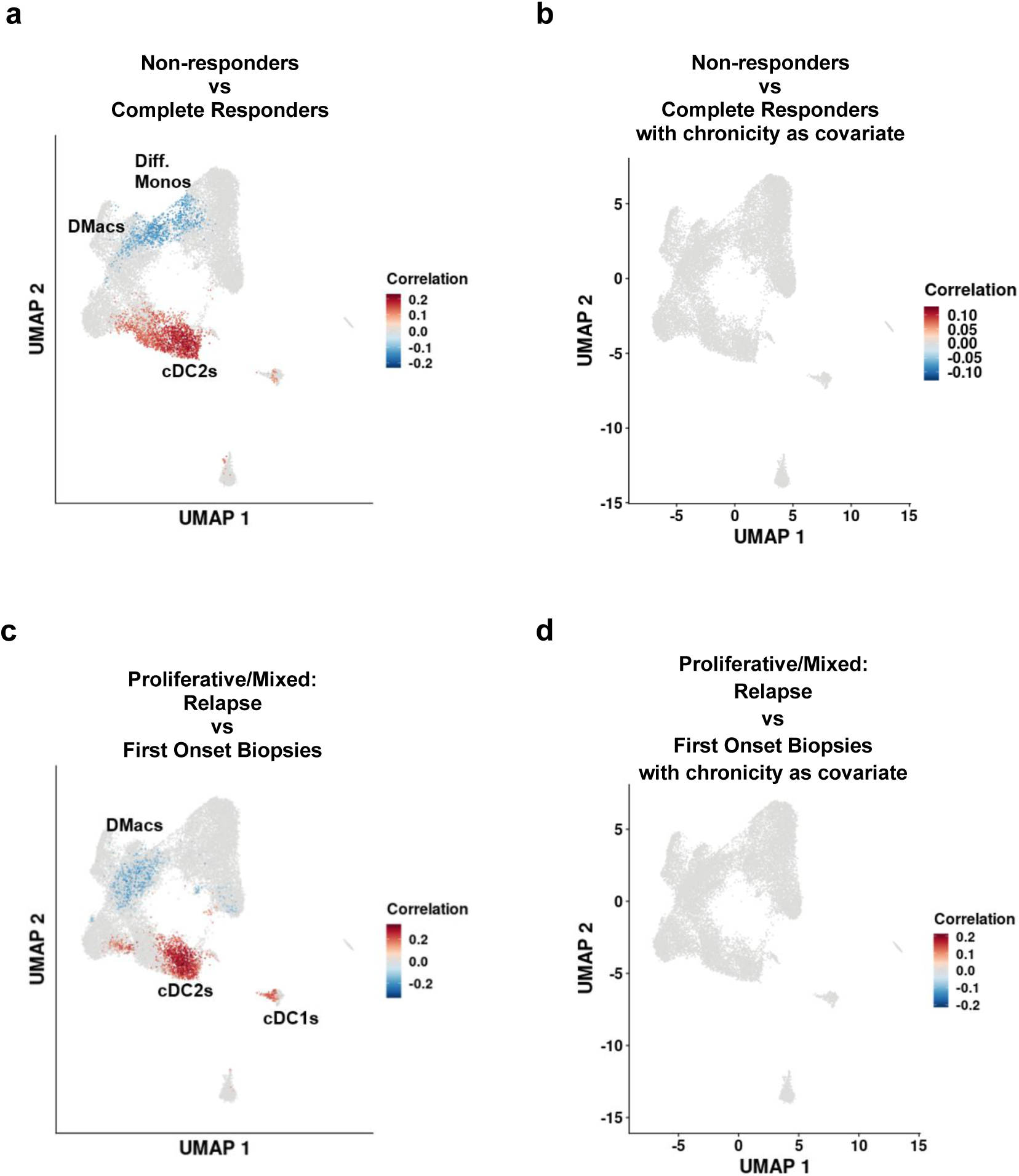
(a) The results of running CNA, comparing non-responders and complete responders, taking all LN patients together. Cell populations with a statistically significant (FDR ≤ 0.1) change in density are colored in red (designating expansion in non-responders) or blue (designating depletion in non-responders). Note that statistically significant correlations were observed only when requiring the UPCR to be less than 0.8 when defining complete clinical response; setting this threshold to lower values did not yield statistically significant correlations. (b) The same as (a), with the CI taken as covariate. No statistically significant correlations were observed with FDR ≤ 0.1. (c) The results of running CNA, comparing first biopsies (generally representing biopsies taken at disease onset) and subsequent biopsies (generally representing biopsies taken at disease relapse), for the proliferative/mixed case. Cell populations with a statistically significant (FDR ≤ 0.1) change in density are colored in red (designating expansion in subsequent biopsies compared to first biopsies) or blue (designating depletion in subsequent biopsies). Note that for pure membranous LN no significant correlations are observed. (d) The same as (c), with the CI taken as a covariate. No statistically significant correlations were observed with FDR ≤ 0.1.

